# Phylogenetic-informed graph deep learning to classify dynamic transmission clusters in infectious disease epidemics

**DOI:** 10.1101/2022.04.10.487587

**Authors:** Chaoyue Sun, Yanjun Li, Simone Marini, Alberto Riva, Dapeng O. Wu, Marco Salemi, Brittany Rife Magalis

## Abstract

In the midst of an outbreak, identification of groups of individuals that represent risk for transmission of the pathogen under investigation is critical to public health efforts. Several approaches exist that utilize the evolutionary information from pathogen genomic data derived from infected individuals to distinguish these groups from the background population, comprised of primarily randomly sampled individuals with undetermined epidemiological linkage. These methods are, however, limited in their ability to characterize the dynamics of these groups, or clusters of transmission. Dynamic transmission patterns within these clusters, whether it be the result of changes at the level of the virus (e.g., infectivity) or host (e.g., vaccination implementation), are critical in strategizing public health interventions, particularly when resources are limited. Phylogenetic trees are widely used not only in the detection of transmission clusters, but the topological shape of the branches within can be useful sources of information regarding the dynamics of the represented population. We evaluate the limitation of existing tree shape statistics when dealing with smaller sub-trees containing transmission clusters and offer instead a phylogeny-based deep learning system –DeepDynaTree– for classification of transmission cluster. Comprehensive experiments carried out on a variety of simulated epidemic growth models indicate that this graph deep learning approach is effective in predicting cluster dynamics (balanced accuracy of 0.826 vs. 0.533 and Brier score of 0.234 vs. 0.466 in independent test set). Our deployment model in DeepDynaTree incorporates a primal-dual graph neural network principle using output from phylogenetic-based cluster identification tools (available from https://github.com/salemilab/DeepDynaTree).

## Introduction

Epidemiological modeling of the spread of a disease during an outbreak is critical in predicting the devastation caused by the root pathogen and the outcome of specific public health interventions. The vast majority of the models used, however, often assume random mixing of the host population, which can affect parameter estimates used in these predictions (e.g., (1–3)). Awareness of this limitation often exists, but incorporation of the numerous relevant structures within a population requires specific *a priori* information regarding not only population behavior but also transmission routes of the pathogen responsible for the outbreak. For example, increased transmission of food-borne pathogens may be found among a community of individuals that primarily rely on a local food vendor (4). Moreover, the structure of the population may be dynamic, as in the case of brief social gatherings wherein air-borne pathogens are more easily spread. Groups of infected individuals for which risk of pathogen transmission is heightened, regardless of their episodic nature, can be identified through combined pathogen molecular data and patient information (e.g., contact tracing). The identification of these groups can provide invaluable insight into specific patterns of spread for which targeted interventions can aid in curbing an outbreak.

Genomic sequence data has become increasingly standard in outbreak surveillance primarily owing to the evolutionary information it provides for the development of therapeutic interventions, a primary recent example being SARS-CoV-2 (5). The evolutionary trajectory of a pathogen can be readily monitored owing to the rapid accumulation of mutations that result from the short generation time and/or infidelity of the replication machinery characteristic of micro-organisms such as viruses and bacteria. The rate of accumulation of mutations, particularly for viruses (6), is often deterministic and so is proportional to the number of replication cycles. Assuming the replication rate is relatively constant for a particular pathogen during infection, the number of replication cycles, and thus mutations, can be estimated from the time of infection of one individual and time of transmission from that individual to another. Fewer mutations are thus expected to occur for shorter transmission times during situations where transmission is more likely. This relationship of evolution and transmission is the principle for genetic clustering, wherein in a sample of a population, transmission clusters are defined as groups of patient-derived pathogen sequences characterized by minimal genetic variation, representing high-risk transmission.

Molecular transmission clusters can be identified using a variety of algorithms that rely on genetic distances. Primarily distance-based methods (7, 8) rely on a user-specified distance threshold, below which samples are considered to be connected. These methods are fast and are accurate; however, the resolution of these methods is limited to the level of the group, as specific mutation information is not used to resolve individual relationships. Alternatively, a phylogenetic tree reconstructed from the set of mutational information at genomic sites offers increased resolution for individual relationships within the group. The collection of relationships between individual samples, represented by branch lengths and branching patterns within the tree, can adopt generalized shapes that can provide additional information as to the underlying contact dynamic (9, 10) (e.g., presence of “superspreaders”) and population dynamic over time (11) (e.g., continued exponential accumulation of infections). In the example of a localized community outbreak, transmission is inherently dynamic over time, typically characterized by early, unchecked growth followed by a natural decline owing to the decrease in number of susceptible individuals (i.e., all individuals are ultimately either infected or immune). The branching patterns within a tree typifying this dynamic are used by Barido-Sottani *et al*. to identify transmission clusters and estimate relevant epidemiological parameters (12), but variation in transmission dynamics over time can present in a variety of flavors, even among community-based clusters, owing to factors such as limited temporal sampling or change in the susceptible fraction of the population. In making the assumption that the number of susceptible individuals is already beginning to decline, one is restricted to *post hoc* outbreak analysis, whereas the identification of clusters of transmission is most critical during the early stages of an outbreak, prior to the observance of this decline and during which design of a more targeted use of often limited public health resources is necessary. Another assumption being made in this model is that the community is isolated, with no migration that would influence the number of susceptible individuals. The susceptible dynamic is perhaps even more relevant for non-community clusters, such as growing sexual networks for which the susceptible population is continually replenished (13).

Despite the importance of both identification and characterization of dynamic transmission clusters in the early recognition of high-risk transmission groups, particularly in the context of public health intervention strategies, no method currently exists to perform these tasks. Various statistics exist to describe the topological tree shape as it applies to population dynamics (e.g., (14–17), also covered in more depth in the Supplementary Material); however, we demonstrate herein that when applied to dynamic transmission clusters, even in combination, they fail to accurately classify these clusters, and further describe the limitations to these generalized tree shape statistics. To address the limitations of existing models, we propose *DeepDynaTree*, a method based on graph neural network (GNN) principles that is capable of learning both the pre-calculated tree statistics and phylogenetic tree structure to accurately predict the dynamics of transmission clusters within the tree. In recent years, the GNN approach has been successfully applied in an increasing number of domains wherein the data are represented as a graph structure connected via complex relationships. Few efforts have been made, however, to exploit the GNN’s capacity to learn from the phylogenetic tree as an existing graph, especially as it is applied to sub-trees, such as transmission clusters. Another unique contribution of this method is an expanded message-passing mechanism from nodes within the tree to edges using the dual graph system, whose vertices and edges correspond to the vertices and edges of the original phylogenetic tree. We thus propose a Primal-Dual Graph Long Short-Term Memory (*PDGLSTM*) model, which alternates to pass the learned state information between the primal and dual graphs. In *DeepDynaTree*’s *PDGLSTM* model, each node and edge of the phylogenetic tree has two states - one hidden, serving as an internal representation of nodes or edges, and the other a memory cell state acting to recall all historic information since the initial update. Two parallel LSTM (18) modules (i.e., Node-LSTM and Edge-LSTM) are utilized to maintain and update the above states. With the iterative node-edge communications, the Primal-Dual Graph design enables learning of the representation of dynamic transmission clusters using the information encoded in the underlying phylogenetic tree edges in a memory-driven fashion.

Using simulated early outbreaks with high-risk clusters, we demonstrate herein the effectiveness of our method and provide detailed interpretation of the model predictions. The consistent superior performances exhibited by *DeepDynaTree* and its variants demonstrate its promising capacity to improve public health efforts in fighting existing and future pandemics.

## Results

### Successful computational workflow for dynamic transmission cluster classification

The overarching goal of this study was to successfully predict the dynamics of transmission clusters based on a subset of previously developed, generalized tree shape statistics, as well as the underlying phylogenetic tree shape. Fig. 1a illustrates the entire computational workflow for dynamic transmission cluster prediction. Briefly, simulation of an early-midway epidemic outbreak was performed using the *nosoi* (19) agent-based stochastic simulation platform, which is designed to take into account the influence of multiple variables on the transmission process (e.g., population structure and dynamics) to create complex epidemiological simulations (see Methods and Table 2). The resulting transmission network is translated in *nosoi* to a strictly bifurcating tree, representative of the underlying pathogen evolution, and used as ground truth in the training and evaluation of the prediction algorithms. Tree nodes were categorized as belonging to transmission among the background (i.e., majority) infected population or to one of seven risk groups comprising clusters of transmission. Risk group nodes were classified into three main categories - static, growing, or decaying - based on predefined transmission dynamics in the simulation (Table 2), and the three categories served as the classification labels during modeling. Various statistics describing the topological tree shape were used to represent each risk group node (see Supplementary Section A), and the branching patterns and branch lengths represent the tree topological information. We grouped the cluster prediction methods into two categories according to the different input information for model development - node-based methods, in which each cluster was treated individually and solely utilized the node features (i.e., the generalized tree shape statistics); the other is our proposed *DeepDynaTree*, wherein a GNN was leveraged to encode a phylogenetic tree as an existing bi-directed graph and directly learned from its structure using a messagepassing mechanism. We also, however, go one step further to propose a novel variant of GNNs, referred to as *PDGLSTM* (Fig. 1b), which enables the incorporation of the phylogenetic topological data and memory-driven processing of tree shape descriptors at nodes and edges.

**Fig. 1.**
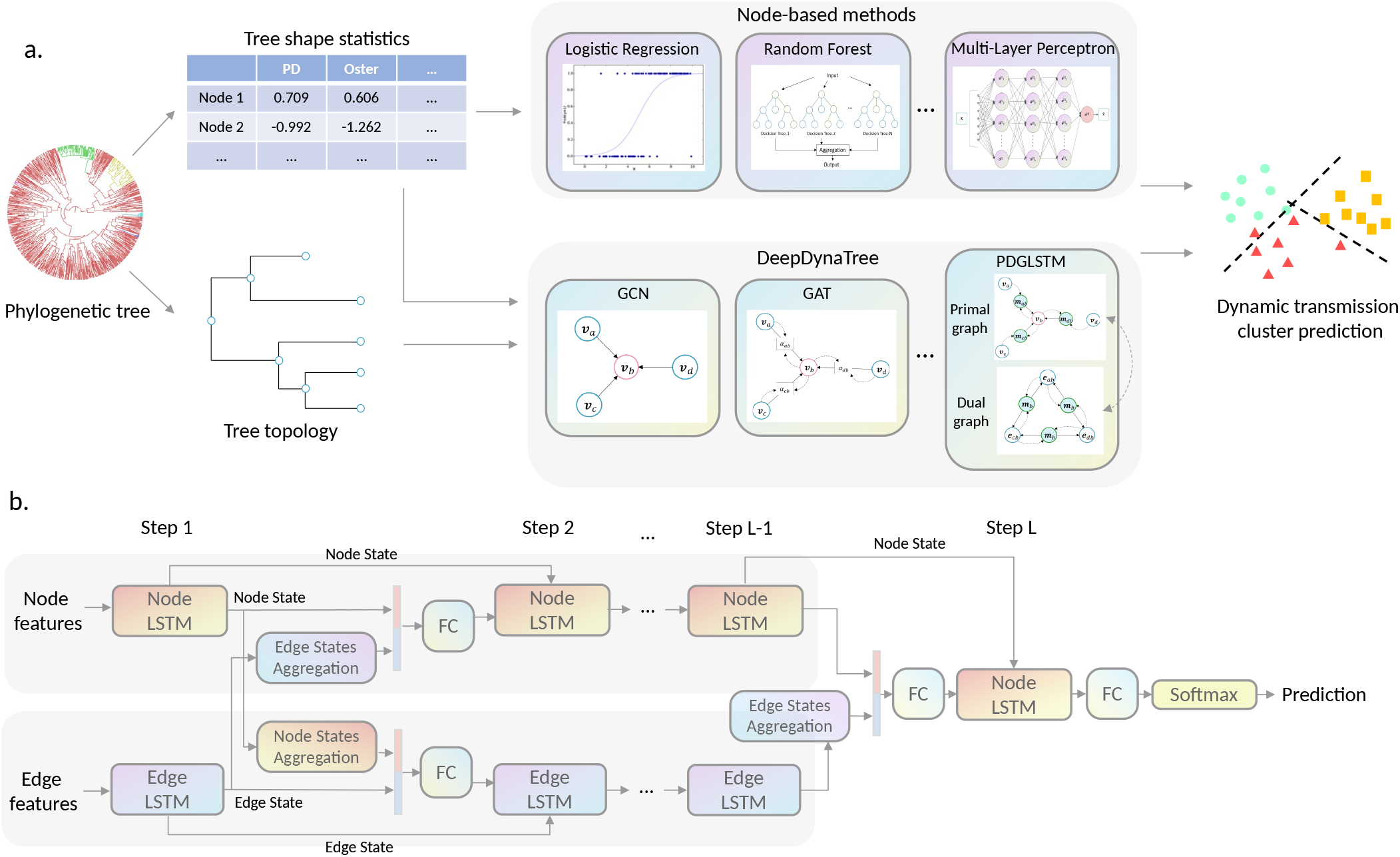
Schematic of dynamic transmission cluster classification workflow and our proposed *DeepDynaTree-PDGLSTM* architecture. **a,** Simulation of an earlymidway epidemic outbreak was performed using the *nosoi* agent-based stochastic simulation platform to create complex epidemiological transmission networks, which were translated to strictly bifurcating, phylogenetic trees. Conventional node-based methods used in dynamic prediction include various machine and deep learning algorithms and only utilize the information in the form of tree shape statistics to predict the dynamics of simulated transmission clusters. By contrast, our proposed graph neural networkbased *DeepDynaTree* method is capable of learning both the statistics and topological structure, including connectivity and branch lengths, leading to significantly better performances. We compare different GNN variants within *DeepDynaTree*, in addition to proposing a new variant referred to as the primal-dual graph long short-term memory (*PDGLSTM*) model, which further boosts the prediction performance by a large margin. **b,** The architecture of our proposed *PDGLSTM*. Two parallel LSTM models take the node and edge features as the initial inputs and produce a set of hidden states. For the node feature updating, an edge state aggregation module is first applied to aggregate the messages from the inbound edges, then the concatenated representation, including the edge messages and the last node features, is passed to the Node-LSTM model to generate the new node state. A similar strategy is also used for edge feature updates. After several rounds of node and edge feature updating and communication, the hidden state of the Node-LSTM is used to predict the corresponding node’s cluster type.

### Method performance and feature importance

This section compares various prediction methods (described in more detial in the Methods), as well as a quantitative evaluation of the importance of different tree shape statistics (described in Supplementary Section A) in dynamic transmission cluster classification tasks. Table 1 demonstrates the detailed comparison of six measurements for the global classification performance on the simulated test set. The metrics including accuracy, F1-score, precision and area under the receiver operator characteristic curve (AUROC), are all equally aggregated over all classes to give the same importance to each class. Brier score (BS) and cross entropy (CE) were used to evaluate the models’ “soft” predictions, wherein the metrics were calculated on the prediction probabilities (see Methods section for details on performance evaluation). *DeepDynaTree* outperformed all node-based methods over all the evaluation metrics by a large margin (see Methods sections for detailed background on prediction methods evaluated). For example, on the two critical classification metrics - balanced accuracy and macro AUROC, *PDGLSTM* achieved 0.826 and 0.934, respectively, and the graph attention network (GAT) achieved 0.703 and 0.860. Arguably, Random Forest (RF) could be considered a reference point of node-based methods because, within the node-based methods, RF achieved the best performance on three of six measurements (including the above two critical metrics) and comparable result with the top performer (XGBoost) for the macro F1 value, whereas MLP and TabNet achieved the best in terms of BS and CE. When compared to RF, even the basic graph convolutional network (GCN) model within *DeepDynaTree* achieved 20.6% and 13.0% relative improvement on balanced accuracy and macro AUROC compared with RF. With *PDGLSTM*, improvement further extended to 55.0% and 25.4% for the above two global metrics, respectively. In terms of the “soft” prediction evaluation, *DeepDynaTree* consistently exhibited remarkable improvement - for example, *PDGLSTM* reduced the best-performing node-based method CE score by more than half (from 0.804 to 0.372), and a similar trend was also observed for BS (from 0.466 to 0.234). Fig. 2a illustrates the macro-averaging ROC curves of the top competitors, where the top three performers all belonged to *DeepDynaTree*’s variants. These results suggest that *DeepDynaTree* shows a distinct improvement in the averaged transmission cluster prediction capacity over the three target cluster types (growing, static, decaying).

**Table 1.** Performance for classification of dynamic transmission clusters in the test data set compared with seven node-based methods, three GNN variants, and three ablation models. The selected node-based models include Logistic Regression (LR), Random Forest (RF), Extreme Gradient Boosting (XGBoost), DeepSet, SetTransformer, Multilayer perceptron (MLP), and TabNet. Under our proposed *DeepDynaTree* framework, the selected GNN variants include Graph Convolutional Network (GCN), Graph Attention Network (GAT), Graph Isomorphism Network (GIN), and the newly designed Primal-Dual Graph Long Short-Term Memory (*PDGLSTM*). Metrics including accuracy, F1-score, precision and AUC are equally averaged over three class types to eliminate the effort of unbalanced label distribution. Brier score (BS) and cross entropy (CE) are used to evaluate the models’ “soft” predictions, wherein the metrics were calculated on the prediction probabilities (see Section for a detailed description). The best performance of each evaluation metric within two different methods is represented in bold.

| Methods | Models | Balanced accuracy ↑ | Macro F1 ↑ | Macro Precision ↑ | Macro AUROC ↑ | BS ↓ | CE ↓ |
| --- | --- | --- | --- | --- | --- | --- | --- |
| Node-Based Models | LR | 0.518 | 0.399 | 0.427 | 0.722 | 0.537 | 0.924 |
|  | RF | **0.533** | 0.396 | **0.436** | **0.745** | 0.514 | 0.894 |
|  | XGBoost | 0.518 | **0.400** | 0.432 | 0.729 | 0.571 | 0.976 |
|  | DeepSet | 0.501 | 0.378 | 0.414 | 0.700 | 0.503 | 0.862 |
|  | SetTransformer | 0.506 | 0.380 | 0.419 | 0.678 | 0.543 | 0.927 |
|  | MLP | 0.523 | 0.387 | 0.427 | 0.618 | 0.503 | 0.852 |
|  | TabNet | 0.525 | 0.393 | 0.429 | 0.661 | **0.466** | **0.804** |
| DeepDynaTree | GCN | 0.643 | 0.542 | 0.546 | 0.842 | 0.351 | 0.598 |
|  | GAT | 0.703 | 0.612 | 0.598 | 0.860 | 0.311 | 0.524 |
|  | GIN | 0.710 | 0.601 | 0.621 | 0.881 | 0.353 | 0.601 |
|  | PDGLSTM | **0.826** | **0.721** | **0.683** | **0.934** | **0.234** | **0.372** |
|  | PDGLSTM w/o [D]<sup>1</sup> | 0.739 | 0.656 | 0.642 | 0.910 | 0.271 | 0.446 |
|  | PDGLSTM w/o [L]<sup>2</sup> | 0.819 | 0.707 | 0.664 | 0.927 | 0.254 | 0.409 |
|  | PDGLSTM w/o [B]<sup>3</sup> | 0.648 | 0.494 | 0.489 | 0.799 | 0.427 | 0.693 |

<sup>1</sup> PDGLSTM without the dual graph design.
<sup>2</sup> PDGLSTM without LSTM module to maintain the historic state information.
<sup>3</sup> PDGLSTM without bidirectional message passing design.

**Fig. 2.**
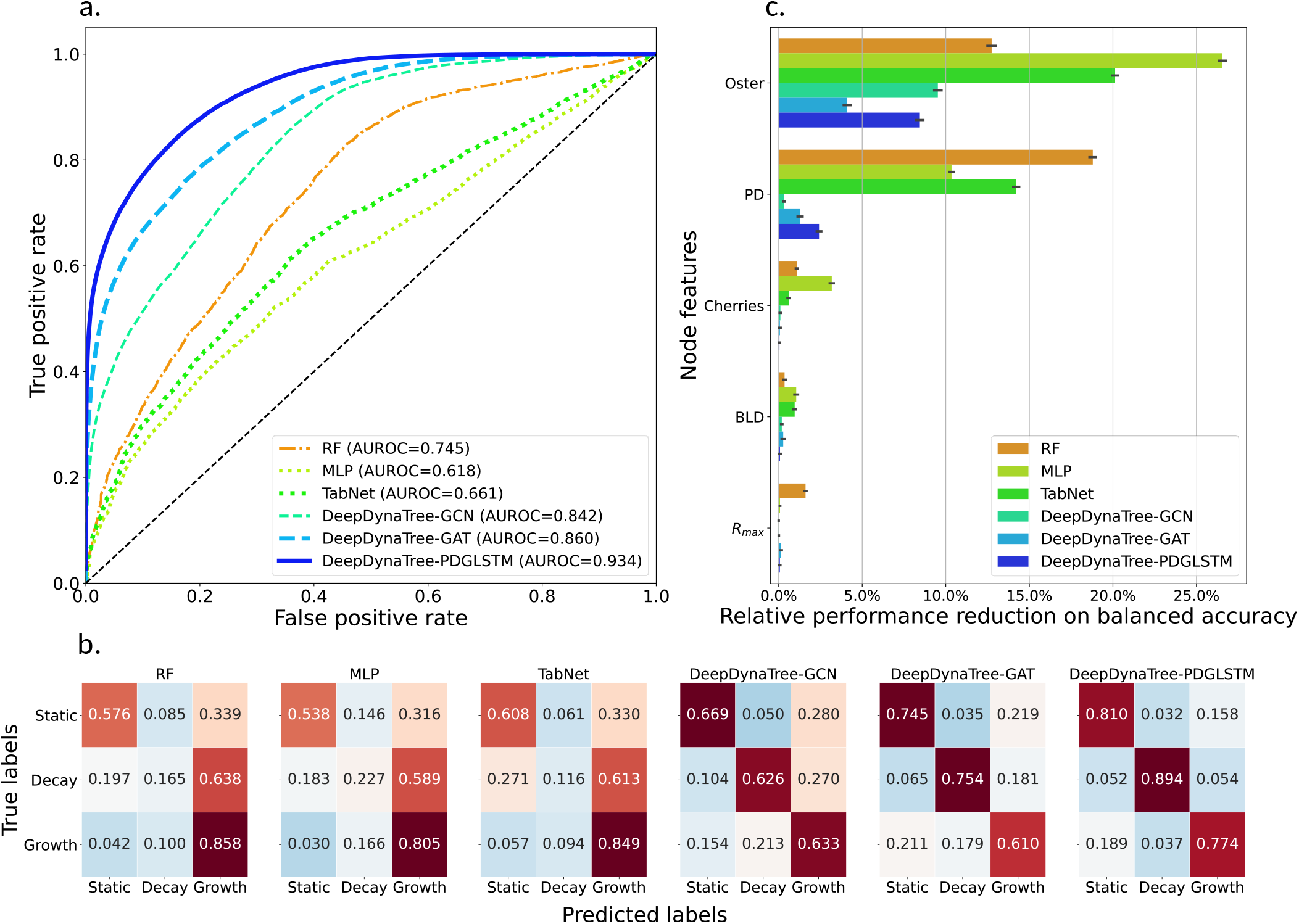
Figurative performance comparison of six selected competitive classifiers and permutation feature importance results. **a,** Comparison of various classifiers, shown on an equally averaged receiver operator characteristic curve (ROC), with AUC indicated for each classifier. **b,** Comparison of various classifiers, shown on confusion matrices, and elements are row-wise normalized by class support size. **c,** Relative permutation feature importance results measured by the balanced accuracy. x-axis represents the relative performance reduction in percentage compared to the non-permutated model and y-axis represents the top-5 most important tree shape statistics features. The features are ranked based on the average relative reduction over all models, and the mean and standard deviation were measured over 50 runs.

Compared with all the current GNN variants shown in the second part of Table 1, *PDGLSTM* achieved the best performance on all evaluation metrics, including the “hard” and “soft” measurements. Taking the second-best model (GAT, comparable with graph isomorphism network [GIN]) as a strong competitor, *PDGLSTM* enhances the balanced accuracy and macro AUROC from 0.703 to 0.826 and 0.860 to 0.934, respectively, and reduces the BS from 0.311 to 0.234. *PDGLSTM* also remarkably outperforms GCN by relative improvements of 28.5% and 10.9% on balanced accuracy and macro AUROC, respectively. Additionally, the confusion matrices shown in the Fig. 2b further demonstrate that the *PDGLSTM* outperforms all GNN competitors on all cluster types. For example, 81.0% of static, 89.4% of decaying, and 77.3% of growing clusters were correctly predicted by *PDGLSTM*, whereas the GAT model correctly identified 74.5%, and 75.4%, and 60.9% of clusters, respectively, and the performance of GCN further dropped to 66.9%, and 62.6% and 63.3%, respectively. These results demonstrate that explicitly framing the time and genetic distance information into the edge-centric dual graph learning process, rather than only utilizing the connectivity of transmission clusters, effectively improves the representation learning capacity of GNNs, leading to more accurate inferences. Fig. 3c, d provide t-SNE plots for the nodes’ latent representations (the input to the last fully connected layer) learned by GAT and *PDGLSTM*. It is clear that the representation of different nodes’ types were better separated for *PDGLSTM*, confirming our design can learn much enhanced representations.

**Fig. 3.**
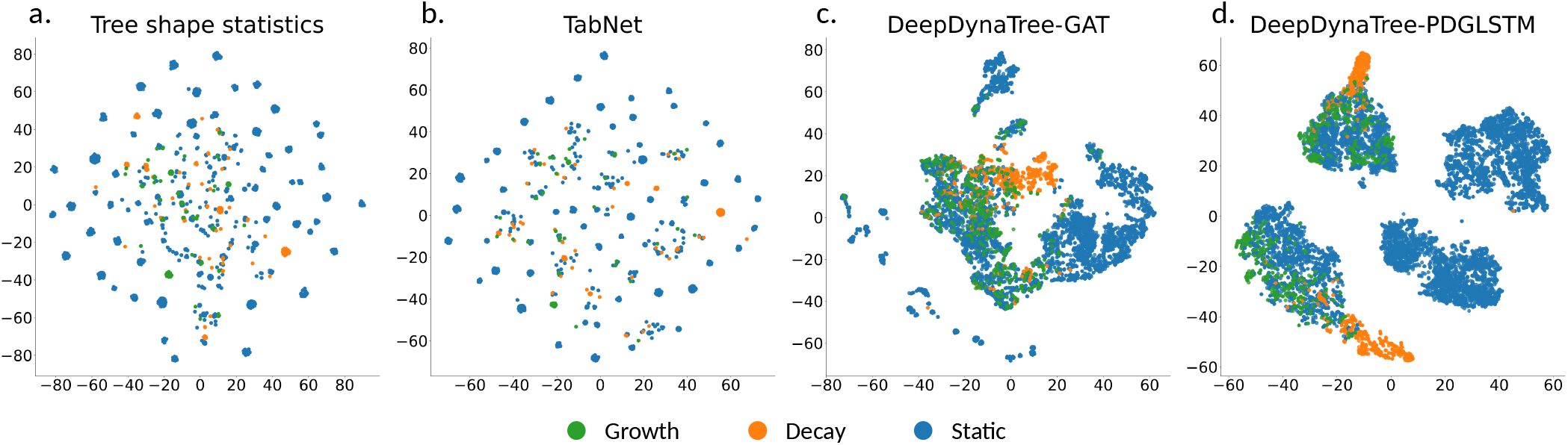
Visualization of the generalized tree shape statistics and the latent representations learned by various methods. **a,** t-SNE plot of the generalized tree shape statistics. **(b, c, d)** are the t-SNE plots for the latent representations of TabNet, GAT and PDGLSTM, which are extracted before the last classification layer. All subplots were generated using the same nodes samples from 50 phylogenetic trees randomly sampled from the test set. The original dimensions were projected to two dimensions with the t-SNE technique.

In addition to comparing methods on their general and averaged performances, we also used the confusion matrices to summarize and illustrate their detailed prediction distributions over different cluster types. Fig. 2b illustrates the confusion matrices of different approaches where each element is normalized by the number of true labels summarized over each row for easy comparison. Similar to the averaged performances, *DeepDynaTree* shows consistently more accurate predictions than node-based methods for each of the individual cluster types. For *DeepDynaTree*’s *PDGLSTM*, 81.0% of static, 89.4% of decaying and 77.3% of growing clusters were correctly predicted, whereas the partitions for the best node-based method (RF) were only 57.7%, 16.2% and 85.7%, respectively. Node-based methods predicted the growing clusters with more accuracy than the other two risk/cluster types, tending to misclassify both the static and decaying clusters as growing, with decaying clusters actually more likely than static clusters to be misclassified as growth - taking the RF algorithm as an example, 63.9% of decaying and 33.9% of static clusters were, respectively, incorrectly forecasted as growth. This result was somewhat counter-intuitive owing, in principle, to the closer overall distribution of static clusters to growing clusters; however, it is at the same time not surprising given the reliance of certain tree shape statistics (*R*_0_ and maximum growth rate) on the early, rapid spread characterizing all decaying clusters (Table 2) and limited ability of other statistics (namely *γ* and *LTTshape*) to classify declining populations. By incorporating individual branch lengths, rather than branch length summaries or metrics, within each cluster (i.e., with GAT or *PDGLSTM*), misclassification of decaying clusters was shown to be much less frequent. Fig. 3a depicts the t-SNE (20) 2-dimensional (2D) feature space for generalized tree shape statistics, where 50 phylogenetic trees were randomly sampled from the test set. We can observe that growth and decay clusters were highly overlapped in many cases, and even in the embedding space of TabNet (Fig. 3b), they cannot be separated well, again demonstrating the limitation of existing tree shape statistics for dynamic transmission cluster classification.

**Table 2.** Simulation information for each risk group.

| Risk group | Contact parameters<sup>1</sup> | Mean P(T)<sup>2</sup> | Mean $R_0$ |
| --- | --- | --- | --- |
| A | $\sim \mathcal{N}(16, 1)$ | 0.015 | 2.2 |
| B | $\sim \mathcal{N}(4, 1)$ | 0.09 | 3.2 |
| C | $\sim \mathcal{N}(4, 1)$ | 0.11 | 3.9 |
| D | $\sim \mathcal{N}(6, 1)$ | 0.09 | 4.9 |
| E | $\sim \mathcal{N}(4, 1)$ | 0.12 | 4.3 |
| F | $N_0 = \sim \mathcal{N}(4, 1),$<br>$K = 10,$<br>$r = \sim \mathcal{N}(-0.04, 0.02)$ | 0.11 | 3.9 |
| G | $N_0 = \sim \mathcal{N}(18, 1),$<br>$K = 20,$<br>$r = \sim \mathcal{N}(0.4, 0.2)$ | 0.11 | 17.8 |

<sup>1</sup> Single values represent mean number of contacts for static clusters.
<sup>2</sup> Probability of transmission

Feature (in this case, tree shape statistic) importance was defined to be the decrements of model performance when a single feature value was randomly shuffled (50 times in this study). Hence, the larger the decremental decrease, the more significant the feature was considered to be to model performance. As demonstrated in both Fig. 2c and Supplementary Fig. 8, the metric (“Oster”) originally used to describe human immunodeficiency virus (HIV) transmission rates (16, 21) and phylogenetic diversity (“PD”) were ranked highest for *PDGLSTM*, as well as the other GNN and node-based models. These values were also highly correlated (ranked correlation coefficients > .90 in Supplementary Fig. 3). The relative feature importance of the Oster statistic (8.44% reduced balance accuracy with permutation) over PD was, however, was nearly three-fold (2.42%), indicating this statistic represents the most reliable of the tree statistics and is unrivaled in distinguishing transmission cluster dynamics among the metrics chosen in this study. Branch length differences (BLD) over time and maximum *N_e_* growth rate (*r_max_*), which have been defined exclusively in this study, outperformed the remaining previously published statistics; however, they contributed primarily to *node-based models*, with minimal contribution to *DeepDynaTree*-specific models and were thus ignored in downstream analysis of model limitations.

#### Model ablation study

Two important designs of our *DeepDynaTree-PDGLSTM* model are 1) iterative primary-dual graph communication and 2) node and edge state updating with temporal information. In order to demonstrate their effectiveness, we performed associated ablation studies. For the first design assessment, we removed the edge-centric dual graph [D] so that the model was restricted to the basic connectivity information and passed the message between nodes only. As shown in the lower section of Table 1, CE increased by around 20% and balanced accuracy dropped by over 10% with this removal. To evaluate the second design, we replaced the parallel LSTM models [L] with the parallel fully connected layers so that the states of node or edge at the next step were only determined by the current states and received messages without any historic information. Cross entropy with this replacement increased by 10% and balanced accuracy was reduced by 2.5% from the original model. The reduced performances confirm the effectiveness of both model designs and suggest edge-centric dual graph was more important than the parallel LSTM approach. Additional metrics (macro F1 scores, precision, recall, AUROC, and BS) revealed similar results (Table 1). In addition, when replacing the bi-directional message passing [B] with the uni-directional passing from tree root to leaves, we observed the balanced accuracy largely reduced from 0.826 to 0.648, which demonstrates that propagating the messages in a bidirectional way between tree nodes is also critical to the model performance. It is worth noting that the other three competing GNN variants all adopt the bidirectional message passing mechanism, the results of which corroborate this conclusion.

#### Model Limitation Analysis

In order to pinpoint the limitations of the *DeepDynaTree-PDGLSTM* model for future application, we calculated the predicting accuracy distribution over ground truth cluster characteristics, including sampling fraction, size, and time span of the cluster (Fig. 4). The non-linear shape of the weighted cross entropy loss (WCEL) across sampling fraction intervals was consistent with the relationship described in the Supplementary Section A, wherein a threshold of sampling is exhibited after which over-sampling also likely influences features based on *N_e_* estimates, approaching lower-reliability estimates as a result of falsely classified clusters as epicenters of transmission. Alternatively, reduced accuracy was observed for small values of cluster size and time span (Fig. 4) - based on the confidence values of WCEL, we could readily conclude that when the cluster size is < 41 individuals and/or the cluster has occurred over a time span of less than 22 days (from first sampled infected individual to most recent), the *PDGLSTM* model is much less reliable. As expected, time span and cluster size exhibited a positive correlation (coefficients > 0.50 in Supplementary Fig. 3 and 4). However, cluster size exhibited a non-linear trend in WCEL, demonstrating that this correlation is limited and that larger clusters with limited time spans (e.g., a burst in transmission) can pose issues for model classification. Time span was also highly correlated with the Oster statistic (coefficients = 0.98), indicating a link between transmission rates and the maintenance of infection among a risk group within a population over time. Exhibiting a similar pattern in WCEL, Oster values < 24 were considered to provide unreliable classifications using the model. Owing to its relationship with time span, however, the calculation of Oster values for clusters prior to assessing confidence in *DeepDynaTree* classification can be accurately replaced with knowledge of the cluster time span.

**Fig. 4.**
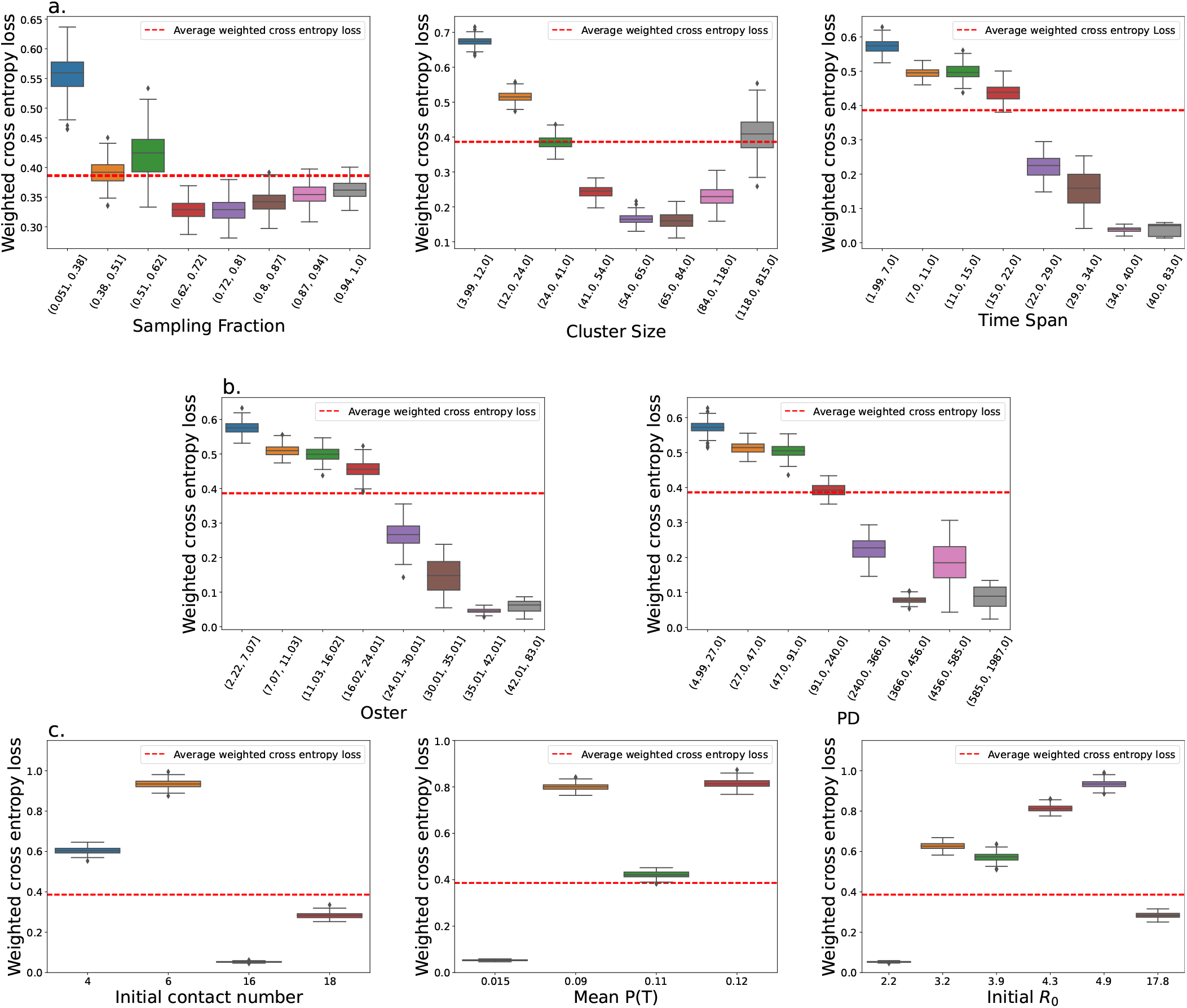
Sensitivity of the *DeepDynaTree PDGLSTM* model to important features and cluster attributes. Each plot shows the weighted cross entropy value for each interval or category. Binned intervals for quantitative data were generated using eight quantiles. Red dashed lines show the weighted cross entropy loss and 95% confidential intervals for the test and provide a threshold for model performance sensitivity. The boundaries of confidential intervals are close to each other so the intervals are unclear. **(a.)** All ground truth cluster attributes. **(b.)** Primary contributing tree statistic features. **(c.)** Varying risk groups within the static category of transmission clusters (see Table 2).

The parameters used to describe transmission characteristics for each risk group in the simulation were similarly evaluated for their impact on prediction accuracy, demonstrating again the slight difficulties in discriminating certain static, growing, and decaying clusters for the *PDGLSTM* model Fig. 4c, as in Fig. 3c and Supplementary Fig. 8, though improved dramatically over remaining models. When evaluated in more depth, lower initial contact numbers (*N*_0_ in equation 1) among individuals within a cluster posed a challenge for discrimination, as did intermediate probabilities of transmission and initial estimates of *R*_0_ (see Table 2). However, this finding was not entirely surprising, as static and growing clusters were allowed to form with similar transmission characteristics, only to differentiate later in time, during which tree topology data are important.

## Discussion

Identification of transmission dynamics among risk groups of pathogen infection is of utmost importance to both basic and translational research alike, providing retrospective understanding of the driving factors of spread and real-time insight into the impact of public health interventions. Whether it be modifications in risk group behavior, the pathogen genome itself, or to the availability of resources, the impacted transmission trajectory can be altered, even within a subset of infected individuals. When traced in the context of patient and environmental metadata (e.g., geographical origins and vaccine availability), these alterations provide critical information that aids in how to combat further spread or future pandemics caused by similar pathogens. Whereas sampled pathogen sequence data, specifically the phylogenetic relationships among these sequences, have been utilized in the estimation of relevant population genetic and/or epidemiological parameters (e.g., *R*_0_), we demonstrate that these estimates are unreliable for smaller datasets, such as individual transmission clusters, even when combined into complex models (e.g., Random Forest). We proposed instead to learn these dynamics jointly from commonly used tree statistics and the underlying tree data by developing a phylogenetic-informed neural network platform – *DeepDynaTree*. We further propose a novel variant of the long short-term memory neural network approach (*PDGLSTM*), which performs iterative message passing between the primal and dual graphs along the phylogenetic tree’s topological structure, utilizing information from both connectivity of the nodes and branch lengths. When integrating these multiple sources of information, the *PDGLSTM* model is capable of generating more informative representation of transmission clusters and enhanced predictions of cluster behavior in terms of growth or decay (or lack thereof).

As with every analytical approach, *DeepDynaTree* is not without its limitations; however, we have carefully evaluated the performance of the *PDGLSTM* model so as to aid users in understanding the boundaries of reliability during application. For example, users should take caution when interpreting results for small transmission clusters (comprised of < 41 individuals) and/or encompassing a brief sampling time span of < 22 days. Whereas the model was not particularly sensitive to the fraction of the cluster being sampled for sequence analysis, sampling was performed randomly rather than using a time- or size-dependent strategy, which may be more realistic of surveillance data and should be taken into account during analysis as well as further training of *DeepDynaTree*. Growing and decaying clusters in these simulations were also modeled using a single, deterministic model of changing population size, whereas the true time-varying transmission dynamics for risk groups may differ depending on stochastic outside factors that influence individual or virus behavior. A broader simulation study encompassing a wider range of time-varying growth models would not only capture a more realistic epidemic, but allow for a more fine-grained transmission prediction, including growth types and rates. It is also important to note that not all clusters included in the final simulation datasets exhibited growth according to their pre-defined model parameters - emergence of clusters shortly prior to simulation termination does not allow for sufficient time for logistic growth or decline to occur. Filtering of clusters via calculation of true growth rates in the future will allow us to determine its influence on the estimated accuracy of the model. Despite these limitations, we describe the first, and thus most extensive, dynamic cluster simulation set thus far, for which *DeepDynaTree* was able to classify transmission dynamics with > 83% accuracy, surpassing the < 54% accuracy achieved with non-neural network models.

*DeepDynaTree* classifies specific groups of individuals as growing, decaying, or static, taking as input *user-specified* transmission clusters, which can be identified using any number of phylogenetic-based cluster-identification algorithms (e.g., (22, 23)) (re-formatting described in https://github.com/salemilab/DeepDynaTree). By doing so, the assumption is made that input clusters have been identified correctly; however, the accuracy of these tools in identifying dynamic clusters has yet to be determined. These approaches are also *themselves* limited in their reliance on thresholds for summary statistics of pairwise genetic distances among samples, which have been demonstrated to differ substantially across transmission pairs and clusters in the study of human immunodeficiency virus (HIV) (8, 24). These differences are influenced by factors such as the extent of clinical follow-up (8) and fraction of the risk group sampled (25–27) and provide the potential for the propagation of error in cluster analysis. Additional training of *DeepDynaTree* for not only characterization, but identification, of dynamic transmission clusters among sampled individuals within a phylogeny may improve our ability to detect clusters of interest and should be explored.

Public health organizations are increasingly utilizing cluster-identification tools in ‘near real-time’ to identify ongoing pathogen outbreaks (26, 28, 29) in order to not only reconstruct the risk factors and aetiology (30, 31) but to prioritize groups for prevention initiatives such as access to preexposure prophylaxis (32). Prioritization is often based on stricter genetic distance thresholds employed in the analytical identification of clusters as described above. For example, whereas a genetic distance threshold of 1.5% is commonly used to define a likely transmission pair for HIV infection (33), the United States Centers for Disease Control and Prevention more recently recommended a threshold of 0.5% (34), representing more rapid transmission and growth of the risk group population. For recently emerging pathogens that quickly accumulate to pandemic status with limited resources for ‘real-time’ epidemiological analysis, optimal genetic distances from known transmission pair sequence data is difficult to achieve, as we have seen with severe acute respiratory coronavirus 2 (“SARS-CoV-2”). Moreover, this approach assumes an unrealistic steady-state system, whereby the more rapid transmission rates experienced by the prioritized risk groups are the result of a risk factor(s) that has/have been present at unvarying levels since initial infection within the cluster. Understanding transmission at a level of resolution that allows us to also identify individuals that have departed from previous risk norms is required for real-time integration of public health and molecular epidemiological data.

Our results confirm that *DeepDynaTree* is a promising tool for transmission cluster characterization that can be modified to address the existing limitations and deficiencies in knowledge regarding the dynamics of transmission trajectories for groups at risk of pathogen infection. Founded on well-developed deep learning techniques, *DeepDynaTree* allows for continual learning of both ground truth and empirical data as they become increasingly available with elevated pathogen sequencing efforts.

## Methods

### Simulation of dynamic transmission clusters

Simulation of an early-midway epidemic outbreak was performed using the *nosoi* (19) agent-based stochastic simulation platform, which is designed take into account the influence of multiple variables on the transmission process (e.g. population structure and dynamics) to create complex epidemiological simulations. Beginning with a single infected individual, rate of transmission of infection to susceptible individuals was dependent on the time since initial infection, duration of infection, number contacts, infection rate, and the risk group (cluster) to which the susceptible individual belongs (Table 2). During the incubation period (4.2 days after initial infection), infected individuals were not permitted to transmit. Individuals were removed from the simulation after an infectious period of 14 days. The number of contacts per infected individual varied according to risk group and time so as to allow for varying dynamics across risk groups. Infection rate was also allowed to vary according to risk group. Seven distinct risk groups [B-G] were allowed to emerge from the background population (A) with a probability of initial infection of 5 × 10^−4^ after at least two background individuals had been infected at the start of the simulation. This rate was used to ensure clusters did not exceed in number the background population. Following initiation, transmission was isolated to the cluster (i.e., probability of zero for infecting an individual in another cluster). The number of contacts for groups A-E were picked from a normal distribution with group-specific means and standard deviation of one (Table 2), representing clusters for which the rate of secondary infection (*R_e_*) remained steady, or static. The number of contacts for F and G, however, were derived from the following function:

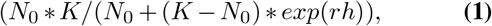

where *N*_0_ is the initial number of contacts, *K* is the maximum number of contacts, and *r* is the rate of change dependent on the current number of actively infected hosts in the simulation (*h*). Cluster F was considered to be experiencing an increasing rate of growth over time, whereas G was considered to be decaying over time (Table 2). Multiple static clusters were also incorporated with varying contact parameters in order to determine indirectly the relationship of these parameters with branching patterns and thus influence on cluster classification. Each of a total of 1,000 simulations was run for 365 days or until a total of 10,000 hosts were infected. One representative clade within the background population was chosen at random from the corresponding internal nodes and maintained all individuals (5-100), representing true clusters of direct transmission. Remaining risk groups were sampled randomly, ranging in frequency from 20-100% of the original cluster population. The background population was also downsampled randomly at a frequency of 20%, representing a more realistic surveillance scenario. Hosts not included within this sample were pruned from the full tree to obtain the final set of simulated trees used for tree statistic calculation and deep learning models. A relaxed molecular clock (evolutionary rate in time) was assumed, and branch lengths were scaled in time (substitutions/site/year) using a uniform distribution rate multiplier 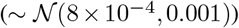, allowing for both genetic distance and time to be used as distinct weights in the neural network models.

In this study, ten thousand simulations beginning with a single infected individual resulted in 9,685 successful outbreaks and thus phylogenetic trees with a grand total of 13,721,009 nodes and 13,711,324 edges. Tree nodes were categorized into two types - nodes belonging to transmission among the background, or majority, infected population (11, 819, 553 nodes in total), and the other belonging to one of eight discrete types of risk groups comprising clusters of transmission (1,901,456 nodes). Risk group nodes were classified into three main categories - static, growing, or decaying - based on pre-defined transmission dynamics in the simulation (Table 2, resulting in 1,547,542 static, 166,723 growing, and 187, 191 decaying samples.

### Input data pre-processing

Because the input distribution was highly unbalanced among the three classification categories, these data represent a realistic scenario in which sampling is biased according to prevalence within the population. Regardless, the data were re-distributed for model development by using a tree-based random split method with a ratio 60%-20%-20%. In other words, all the nodes of a single phylogenetic tree were only allowed to be split into either the training, validation, or test subset with the proposed probabilities, thereby avoiding data leakage, which results from the tendency of nodes (or group of nodes) from a single tree to exhibit high similarity - random splitting of these nodes can produce training data that contain information about the test set, but similar data will not be available in the production environment, resulting in poor generalization capacity. Supplementary Fig. 1, 5 and 6 respectively illustrate the detailed distribution of node, edge, and tree features used in different subsets for the model development and the success of the strategy in providing a more consistent data distribution over different subsets. It is important to note that in our experiments, 5-fold cross-validation was used for all machine learning based methods to select the optimal parameters. Considering the high computational cost introduced by deep learning, we have optimized all deep learning based methods on the above fixed validation data set, instead of using the 5-fold cross validation. Before feeding the data into the learning algorithms, we performed some standard data pre-processing for raw inputs, including quantile-based data discretization, one-hot encoding, and z-score normalization, based on different feature formats. See Supplementary Section B for details. The resulting node and edge features have 16 and 2 dimensions respectively. Supplementary Fig. 2 and 5 provide an illustration for the node and edge feature distribution.

### Potential influence of inherent epidemiological attributes on model performance

Though the metrics used to describe tree shape (see description in Supplementary Section A) are potentially useful in classifying dynamic transmission clusters, each also has the potential to suffer from the reliance on the informativeness of the data, as with other phylogenetic and epidemiological approaches (35). Three attributes describing data informativeness have been incorporated into the models described below in order to evaluate the sensitivity of the models, as well as to allow the user to provide a measure of confidence in the results, given *a priori* knowledge of the data at hand. Sample size is the most basic of these attributes that can wreak havoc on quantitative epidemiological estimates, with smaller sampling sizes providing less reliable estimates owing to the limited amount of information represented. Additionally, considering a large fraction of the tree statistics rely on branch lengths scaled in time, the total time during which a cluster has taken place (calculated using branch lengths) may similarly influence the uncertainty of the estimate. Whereas some level of correlation is expected between cluster size, time span, and true cluster dynamics (e.g., growing clusters should intuitively be expected to be larger in size as well as time span), this relationship breaks down under two scenarios - firstly, a cluster characterized by a greater transmission potential might not emerge until the end of the simulation, which is terminated conditionally on the total number of individuals infected (10,000) or total days occurring (365). The end of the simulation can also be thought of as the most recent collection date prior to data analysis in a real-life study. Secondly, a small cluster size may actually represent a large cluster sampled at low frequency (included in the simulation), as is often a realistic scenario in epidemiology, owing to under-represented populations in health care or undetected transmission among asymptomatic individuals, for example. Hence, if the performance of *DeepDynaTree* were to be heavily influenced by sampling frequency, increased efforts in field investigation would lend increasing confidence in the classification of a cluster. Sampling fraction also has another potential influence on the use of tree statistics in predicting dynamics, as the estimate of *N_e_* described above assumes a relatively small fraction of the total population has been sampled. As this fraction approximates the census, or total population size, external branch lengths are exaggerated, and the tree gives the appearance of inflated growth (36). Hence, while sufficient, unbiased sampling can increase the sample size and thus informativeness of the data, over-sampling (as with a fully sampled transmission chain) may result in falsely classified clusters as epicenters of transmission.

### Phylogenetic tree representations and notations

We represent a phylogenetic tree using a tree structured graph notation *G* = (*V,E*), where *V* = {*v*_1_, …, *v_N_*} is a set of *N* nodes, and *e_ij_* ∈ *E* is an edge from node *v_i_* to node *v_j_*. **v***_i_* and **e***_ij_* denote the features of node *v_i_* and edge *e_ij_*. An adjacency matrix denoted as 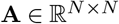 represents the connectivity of various nodes, and the neighborhood set of the node *v_i_* is represented as 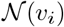. The key annotations are listed in Table 3, where bold uppercase characters are used to denote the matrices and bold lowercase characters denote the vectors.

**Table 3.** Commonly used notations in this paper.

| Notations | Descriptions |
| --- | --- |
| $\mathbb{R}^F$ | $F$-dimensional Euclidean space |
| $G = (V, E)$ | A graph |
| $V = \{v_1, \dots, v_N\}$ | The set of $N$ nodes |
| $e_{ij} \in E$ | An edge $e_{ij}$ from node $v_i$ to $v_j$ |
| $\mathbf{v}_i$ | The features of node $v_i$ |
| $\mathbf{e}_{ij}$ | The features of edge $e_{ij}$ |
| $\mathbf{A}$ | The adjacency matrix |
| $\mathcal{N}(v_i)$ | The neighbors of $v_i$ |
| $\mathbf{W}, \mathbf{U}, \mathbf{b}$ | The learnable parameters |
| $\sigma(\cdot)$ | The activation function |
| $[\cdot, \cdot]$ | The concatenation of vectors |

### Node-based methods using generalized tree shape statistics

We first attempted to assess the value of a subset of relevant tree shape statistics (described in more detail in the Supplementary Section A) in predicting the dynamics of simulated transmission clusters belonging to differing risk groups. Each node within a cluster was represented by 10 such features, and the full data set was represented in a common tabular data format, comprising nodes (rows) with the same set of features (columns). Different machine and deep learning algorithms were developed to predict the nodes’ dynamic characteristic label (static, growing, or decaying), heretofore referred to as the node-based methods because they focus on node features only. The following two subsections describe the developed algorithms in details.

#### Machine learning models for node-based methods

We investigated three broadly-used machine learning methods ranging from logistic regression (LR) (37) to two strong and robust ensemble methods - random forest (RF) (38) and Extreme Gradient Boosting (XGboost) (39). LR served as the simplest baseline method, which fits a generalized linear model with 60 = 10 × 3 × 2 parameters in our experiment to minimize the residual sum of squares between node labels and the linear approximations. To deal with the multiclass case, a multinomial logistic regression was utilized to train the model, and a L2 regularization term with factor 0.001 was applied to prevent over-fitting. RF is a strong and robust bagging ensemble method, which develops diverse decision trees on the bootstrap-sampled subsets of the original training set and then aggregates their outputs as a final prediction (38). In our experiment, we assembled 100 tree classifiers, and in order to control over-fitting, the max depth of each tree was constrained as 5, and the minimum number of samples required to be at a leaf node was set as 14. Another utilized machine learning algorithm, XGBoost, is one of the most popular and effective Gradient Boosting Decision Tree (GBDT) algorithms. Unlike RF, XGBoost utilizes the boosting ensemble method, which fits multiple base models sequentially such that the training of the current base model at a given step depends on how the previous base models fit the training set (39). We selected gradient-boosted decision trees as the base model, and in order to again avoid over-fitting, we slowed the boost learning step by a shrinkage factor 0.3, and the training samples were sub-sampled by 7.79% in every boosting iteration. To reduce the model complexity, the weight for L2 regularization was set as 1, and the depth of trees was constrained to be 20. Both RF and XGBoost models have been successfully applied in many machine learning tasks and achieved state-of-the-art results, especially for input data with tabular format (40, 41).

#### Deep learning models for node-based methods

We also investigated several deep learning-based approaches (i.e., multilayer perceptron (MLP) (42), DeepSet (43), SetTransformer (44), and TabNet (45)). A frequent and straightforward choice of neural network architecture to learn from a set of independent features, MLP was composed in this study of 5 fully connected layers, containing 64 – 64 – 32 – 10 – 3 hidden units. The Rectified Linear Unit (ReLU) was then used as the nonlinear active function. A more complex deep learning architecture for tabular learning, TabNet uses a sequential attention mechanism to select a subset of features to process at each decision step, enabling better interpretability and learning capacity. For TabNet, the dimensions of both prediction layer and attention layer were set to 64. The number of successive steps was set to 10, and each step included 2 independent Gated Linear Unit (GLU) layers and 2 shared GLU layers. The scaling factor for attention updates was set to 2, and momentum for batch normalization 0.02. The masking function sparsemax (45) was used for feature selection.

Besides treating the nodes’ features as tabular data, we also explored another view of the feature format, which considers them as set-structured data. Unlike the fixed dimensional vectors, the output label for the entire set should be invariant to the permutation of set elements (that is, the order of input features). Such problems are widespread, such as 3-dimensional shape recognition from point clouds, wherein the shape label is invariant to the order of points. The models used to address them should be *permutation invariant*, which means the predictions should not depend on the order of elements in the feature set. DeepSets (43) is one of the pioneering works used in solving set-input problems. In the model, each feature of the set is first individually embedded by a neural network, and then a permutation invariant operation (e.g., sum) is applied to aggregate all embeddings, or output of inner layers in the neural network. The final output is generated by applying another neural network on the aggregation in the same manner as in any deep network (e.g., fully connected layers, non-linear active function, etc.). In our setting, both encoder and decoder included 4 fully connected layers and each layer included 256 hidden units. The permutation invariant operation was the average over embedded features of encoder’s output. SetTransformers (44) is another popular permutation invariant method, which improves on DeepSets by using a self-attention mechanism to process every feature in the set. SetTransformers enables discovery and modeling of the potential interactions among each node’s statistic features. In our experiment, the encoder included 3 Induced Set Attention Blocks (ISAB) modules with 2 heads and 64 hidden units. The decoder consisted of 1 Pooling by Multihead Attention (PMA) module and 2 Set Attention Blocks (SAB) with 4 heads.

### *DeepDynaTree* - A phylogenetic-informed approach

Unlike the previously mentioned approaches, our designed *DeepDynaTree* platform is phylogenetically informed, which means it not only utilizes the tree shape statistics used in the node-based models described above but also the underlying topological and branch length information within the tree itself. To achieve this, we proposed the use of a GNN to learn from the raw phylogenetic tree directly and provide the nodewise classification. GNN is a type of deep learning model designed for addressing graph-related tasks. A typical GNN usually consists of several trainable layers and operations (e.g., propagation and aggregation, updating, readout or pooling, etc.) The propagation operation is used to communicate information between nodes or edges based on an information diffusion mechanism so that the aggregated information can capture both nodes’ or edges’ features and phylogenetic information. Then GNN updates nodes’ states by considering the aggregated neighborhoods’ information, or connected edges’, information recurrently. Similarly, the edges’ feature can also be updated by exchanging the connected nodes’ information. When the subgraph’s or graph’s high-level representation is required, a readout operation is used to extract information from nodes and edges, and common techniques for readout, including sum, average and max pooling. These layers are usually stacked repeatedly, and, with the increase of the number of layers, the receptive field size of a node will correspondingly increase, resulting in more high-level and informative representations. For a more comprehensive picture of deep learning on graphs, readers are referred to (46–49). In *DeepDynaTree*, a phylogenetic tree is considered a static bidirectional homogeneous graph, wherein each internal node represents the most recent common ancestor of the two lineages descended from that node and an edge between two nodes represents their evolutionary and/or temporal distance. The dynamic prediction of transmission clusters is formulated as a node-level classification task on the graph. In other words, each risk group node is classified as belonging to the category of static, decay or growth. It is worth noting that we also explored to formulate a phylogenetic tree as a directed graph, which allows to propagate the messages in one direction only, i.e., downwards from tree root to leaves. However, the performance was not satisfied compared with the bidirectional passing, which was demonstrated in the Result Section.

We first assessed three popular GNN variants - Graph Convolutional Network (GCN) (50), Graph Attention Network (GAT) (51), and Graph Isomorphism Network (GIN) (52). In contrast with the node-based approaches, these methods consider the underlying relationship between different cluster nodes, instead of viewing each node individually. However, it is worth noting that these GNN variants can only utilize the basic topological information in the phylogenetic tree (i.e., connectivity) without adequately utilizing the branch length information, representing the genetic distance and/or evolutionary time separating individual nodes within the tree. For example, for GCN, the weights for aggregating neighbors are defined by the tree topological structure (i.e., an adjacency matrix). Similarly, in GAT, the attention coefficient is calculated based on the neighbors’ features and is masked as zero when there exists no direct connection between any two nodes.

GCN is a typical spectral network, which utilizes a first-order approximation of spectral graph convolution with localized filters on the nodes of an input graph (50). Formally, a layerwise propagation rule is defined as:

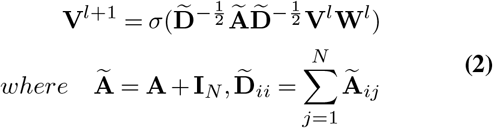

where 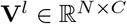 and 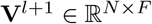 respectively represent the input and output node features for *l*-th layer with *C* and *F* dimensions. **V**^0^ corresponds the original generalized tree shape statistics of all *N* nodes. 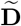 is a diagonal matrix of node degrees and **I**_*N*_ is an identity matrix.

We stacked 11 layers of the spectral graph convolution, and each layer contained 64 hidden units. Unlike the original semi-supervised training scheme proposed in (50), in our experimental setting, we adopted the supervised training manner wherein all the risk group nodes in the test set did not have any label information, and nodes in the training and validation subsets were all labeled.

The second model, GAT, was originally proposed by Velickovic *et al*. (51), which introduces the self-attention mechanism into the graph learning, by which the node state updates depend on the attention coefficients over its neighbors. Specifically, the attention coefficients are calculated as:

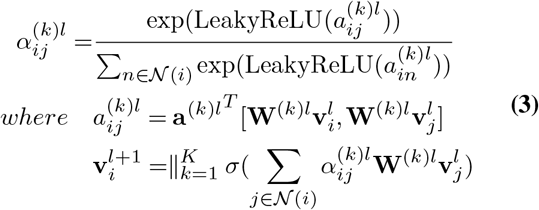

where 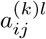 is a scalar representing the importance of node *v_j_* to node *v_i_* for *l*-th layer and *k*-th head, and the attention coefficient 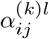 regarding to node *v_i_* is normalized with LeakyReLU non-linear function and softmax function. 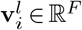 represents the features of node *v_i_*, where *F* = *K* * *H* for *K* heads and *H* output dimensions of each head, and 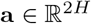 is learnable weight vector. || here stands for the operation concatenating multiple vectors. In our setting, we stacked 8 graph attention layers, the multi-head attention of each layer was applied with the number of 3, and each head contained 64 hidden units.

Xu et al. (52) proposed the GIN variant, which supposedly achieves the maximum discriminative power among graph neural networks. It uses a multilayer perceptron (MLP) model to update the node features as:

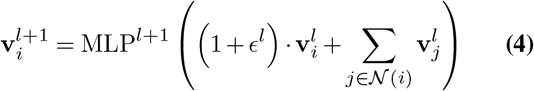

where *ϵ* is either a learnable parameter or a fixed scalar. Different from generating whole graph embedding as proposed in (52), here we directly apply fully-connected layers on nodes’ feaures in each layer and generate the final prediction score by summation:

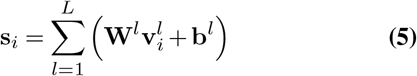

where 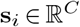 represents the predicted score of *C* classes. *L* is the number of GIN layers. In our implementation, *ϵ* was set as a learnable parameter. we stacked 10 GIN layers, the MLP of each layer was applied with 5 layers, and each layer contained 64 hidden units.

### *PDGLSTM* - An improved GNN variant for accurate dynamic transmission cluster prediction

To overcome the above limitations, inspired by (53, 54), we proposed a Primal-Dual Graph Long Short-Term Memory (*PDGLSTM*) model. The original phylogenetic tree is referred as the *primal graph*. Its dual graph (also known as line, or adjoint, graph in graph theory) was constructed by considering each primal edge as a dual vertex, and two dual vertices considered adjacent if they shared a common endpoint in the primary graph. Supplementary Fig. 9 provides an illustration of the dual graph construction given an example phylogenetic tree. Following this process, *PDGLSTM* generates two disjoint sub-graphs for a single input tree, including a node-centric primal graph and edge-centric dual graph. In order to learn the representations for both nodes and edges, two parallel LSTM modules are employed, respectively, on the primal and dual graphs. With inherent cell and hidden states, this approach enables memorization of the long historic information and the transfer of the temporal signals throughout the process of sequential events. As shown in Fig. 1b, Node-LSTM and Edge-LSTM take the pre-processed nodes and edge features as the initial input. Subsequently, a message passing operation between primal and dual graphs is performed to exchange the information of nodes and edges, wherein the output of a node’s LSTM is passed to its linked edges, and the node itself correspondingly receives the linked edges’s LSTM output as well. Since the maximum degree of a node on the phylogenetic tree is three, during the message passing process, one node can receive at most three edges’ information, while one edge always receives messages from two nodes. All the received messages of a node are first combined by an element-wise addition operation, concatenated with its own previous hidden representation, and fed into a fully connected layer followed by a non-linear activation function to generate the new input for Node-LSTM. As the messages for an edge is always from two adjacent nodes, they are directly concatenated with the edge’s own hidden representation as the input of Edge-LSTM. Integrating the current input with the short-term (hidden state) and long-term (cell state) memories, LSTM is capable of generating a set of new hidden and cell states, which efficiently combines the contextual information from neighbor edges (or nodes) and historic information of the node (or edge) itself. By repeating the message passing and updating steps iteratively, every node and edge can gather the updated states from multi-hop neighborhoods, resulting in enhanced and more informative hidden representation. The current state of each node and edge is thus represented by the hidden state of the corresponding LSTM. In the last iteration step, only node states are updated by Node-LSTM, and the state of each node is used to predict the node type through a fully connected layer followed by a softmax activation, which generates the normalized probability for each cluster type.

For all the deep learning-based methods, the input features were scaled to the same range of [−1, 1] prior to being fed into the network, and a weighted cross-entropy (WCE) loss with inversely proportional to class frequencies weights was applied for model optimization. Hyper-parameters of all the methods were extensively optimized on the same validation set for a fair comparison. Refer to Supplementary Section B for more thorough feature pre-processing, hyper-parameter selection, and training details.

#### Primal-dual state updating

Our proposed *PDGLSTM* formulates the graph representation learning using two disjoint but iterative sub-graphs i.e., node-centric primal graph and edge-centric dual graph, and updates the node and edge embedding using two parallel LSTM models. LSTM (18) model, equipped with the chain-like network structure, is well-suited to extract the temporal sequence information. In the primary graph, each node LSTM takes its aggregated messages as current input and generates the updated node state by jointly considering the current the historic information. Formally, given the hidden state 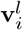 of node *v_i_* at time step *l* and its aggregated message **m**_*i*_, the node state is updated following the *LSTM_node_* updating rule:

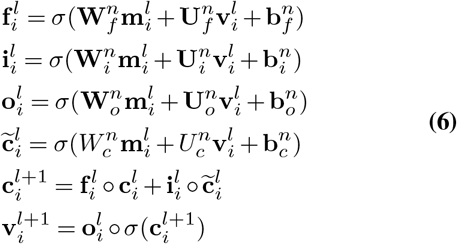

where **W**^*n*^, **U**^*n*^, 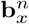 are model parameters for *x* ∈ {*f, i, o, c*}. 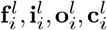 denote the output of forget gate, input gate, output gate and memory cell respectively. Here, the time step *l* represents the *l*-th primary-dual graph communication and it is conceptually similar to the layer index *l* used by other GNN variants.

Similarly, in the edge-centric dual graph, the edge state 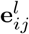 at time step *l* is also updated using a LSTM model:

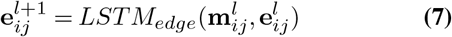

where the **m***_ij_* represents the aggregated message for edge *e_ij_* and the detailed computation of *LSTM* is as same as Eq. 6, except replacing the node inputs with the edge and its associated message information. In our setting, the hidden sizes of both Node- and Edge-LSTM models are configured as 64.

#### Primal-dual message passing

Primal-dual message passing strategy is designed to aggregate and process the corresponding messages for nodes and edges, allowing for information fusing between the node set *V* and the edge set *E*. Specifically, for node *v_i_*, its aggregated message **m**_*i*_ is computed from the current states of inbound edges and node itself:

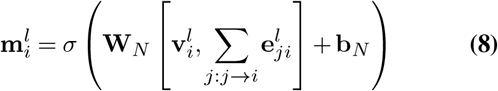

where, *j* → *i* denotes all the inbound edges to the node *v_i_*. Similarly, the message information **m***_ij_* for edge *e_ij_* is generated by:

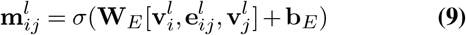

where *σ* here represents the Sigmoid activation function and both **W** and **b** are learnable parameters.

#### Transmission cluster prediction

With iteratively applying the primal-dual message passing and state updating steps, *PDGLSTM* is able to gather extensive information from multi-hop neighboring nodes and edges, resulting in an enhanced and more informative node representation for final prediction. In our setting, the above two steps were repeated 13 times, which means the receptive field of each node was up to 13-hop neighboring nodes and 13 edges. A prediction head with one fully connection layer was utilized on the final high-level node embedding.

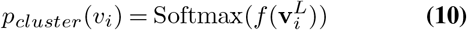

where *f* is realized by a single-layer feed-forward network, and *L* is the total time steps.

### Model evaluation metrics

In this study, our proposed DeepDynaTree was comprehensively evaluated on six performance metrics including macro-averaged precision, F1-score, and area under the receiver operator characteristic curve (AUROC), balanced accuracy, Brier score, and cross entropy. Precision, F1-score and AUROC are broadly used measures of the discriminability of a pair of classes. In order to deal with the multi-class case, we first generated the metric values for each class in the one-vs-rest manner, and then averaged them with equal weights to give the same importance to each class. Considering the extremely unbalanced label distribution in our data set, equally averaging the metrics over all classes instead of weighted averaging by the reference class’s prevalence in the data can effectively avoid overestimating the models that only perform well on the common classes while performing pooling on the rare classes, and this equally averaging strategy is also named as macro-average. The balanced accuracy is defined as the average of recall obtained on each class (55). Besides the above metrics calculated based on the “hard” classification results (i.e., final category information only), we also measured the Brier score and cross entropy to evaluate the models’ “soft” predictions, wherein the metrics were calculated on the prediction probabilities. Eq. 11 illustrates the definition of Brier score (BS) and cross entropy (CE) values,

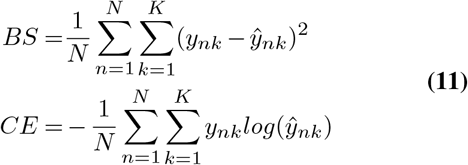

where 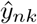 denotes the predicted probability of sample *n* for label *k*, and *y_nk_* is the ground truth with binary value. *N* is the number of samples in the data set and *K* is the number of classes where it equals to 3 in our case. Lower BS and CE values thus indicate better prediction performance.

### Implementation Details

This section introduces the implementation details of our final *DeepDynaTree-PDGLSTM*. The model took approximately 3 days to train for around 400 epochs with mini-batch of size 32. An adam optimizer was used for training with an initial learning rate 3 × 10^−3^, and 90% reduction was applied to the learning rate if the validation loss did not improve in the consecutive 300 epoches until reaching the the minimum value 1 × 10^−6^. We finetuned all the hyper-parameters on the validation data set. All experiments were performed on a workstation with 12 Intel Core i7-5930K CPUs and a single Nvidia GeForce GTX TITAN X GPU card.

## Data availability

Source code used to simulate the data (including seeds) used in this study, as well as code used in calculating tree shape statistics and transforming annotated trees for *DeepDynaTree* classification, are provided in https://github.com/salemilab/DeepDynaTree and are written in R (56).

## Code availability

The code used to generate the results shown in this study is available under an MIT Licence in https://github.com/salemilab/DeepDynaTree.

## ACKNOWLEDGEMENTS

The authors would like to acknowledge University of Florida Research Computing for providing computational resources and support that have contributed to the research results reported in this publication. URL: http://researchcomputing.ufl.edu.

## Supplementary Material

### A. Generalized tree shape statistics evaluated in prediction of cluster dynamics

The information contained in the shape of a molecular phylogeny can be crudely partitioned into two components - the topology (branching structure) and branch lengths (also sometimes referred to as node heights). Although these components are intimately linked, the distinction is nevertheless useful (57). Some tests of evolutionary hypotheses require only tree topology, such as statistics that measure tree balance (17), whereas other tests, particularly those based on the birth-death or coalescent processes (58, 59), depend only on branch length information. The common feature of these tests is that they can be used to estimate the rate (or change of rate through time) of evolutionary processes. For example, when assuming a clock-like evolutionary behavior (i.e., the rate of accumulation of mutations remains constant over time), branch lengths representing genetic differences between samples taken at a specific point in time can be re-scaled in similar units of time, revealing divergence dates, and thus divergence rates over time, among lineages within the tree. Nee *et al*. (59) proposed the lineage through time (LTT) plot as a way to graphically investigate the demographic history of a population using a sample of gene sequences and to estimate birth and death rates in order to test the null hypothesis of a static population. Pybus and Harvey soon after observed that this constant-rates model could be rejected for a given phylogeny if the internal nodes, or divergence events, were closer to the root of the phylogeny than would be expected under a pure birth model (virtually no extinction of lineages) (14). This observation was used to develop a statistic (*γ*) that described the relative position of the nodes within the tree and was used in this study to quantify transmission dynamics. However, it is important to note that the gamma statistic assumes equidistant root-to-tip branch lengths (i.e., all samples were taken at the same time), which is violated in our simulations and in modern epidemic datasets. An additional statistic unbiased by sampling time was thus included (“BLD”), which was calculated as a function of the difference between branch lengths (median) observed in the first and second halves of the time at which the cluster was observed.

Pybus later improved on the LTT plot to provide a framework for inference of the population demographic history (60), which has since undergone several changes, including the implementation of a Bayesian approach employing prior distributions for parameters pertaining to the underlying evolutionary process, as well as the change in effective population size (*N_e_*) (61) in order to account for uncertainty in tree reconstruction. The original “skyline” family of Bayesian phylogenetic frameworks have assumed that *N_e_* follows a stochastic process such as a Brownian motion (60, 62), which was demonstrated by Volz and Didelot (15) to have a potentially large impact on estimates, especially when genealogical data are sparse and uninformative. Volz and Didelot (15) thus recently proposed a modified approach defined according to a growth rate prior that was able to reproduce dynamics estimates (e.g., the initial rate of secondary infection [*R*_0_]) expected under a tested variety of epidemic situations, including otherwise erroneously predicted stable populations. For the simulations described herein, Volz and Didelot’s approach, referred to as *skygrowth*, was used to estimate the effective population size (*N_e_*) for individual risk groups within the tree in order to derive point estimates of potentially relevant epidemic parameters, including *R*_0_, rate of overall *N_e_* growth (“AbsGrowthRate”), maximum rate of growth given discrete intervals of time (“*R_max_*”), and fraction of time spent in the maximum growth phase (“FractionTimeGrowth”) (Table 1).

Branch lengths scaled in time have also been used in other ways to characterize epidemic dynamics. Recently, Oster *et al*. demonstrated that human immunodeficiency virus (HIV) transmission rates for a given cluster (number of transmissions per 100 HIV-infected persons per year) could be estimated as a function of the size in tips, the sum of the branch lengths, and the length of the longest branch in the cluster (16, 21). While only described in the context of a snapshot in time, we aimed to characterize the capability of the “Oster” statistic to distinguish the three classifications of transmission clusters described above (static, growing, and decaying). Testing the hypothesis that this statistic is primarily driven by the sum of the branch lengths, or phylogenetic diversity (PD), this value was also included in the model.

In terms of tree topology, the degree of balance, or symmetry, within the tree has been linked to phylodynamic inference, as it is thought to be influenced by biological factors such as differences in infectiousness or contact rate (63). One measure of imbalance relies on the simple identification of a branching pattern referred to as a cherry. Cherries are defined as two tips that share, or coalesce via, a *direct* ancestor within the tree. The expected number of cherries in a tree with *n* taxa under a pure birth (Yule) model is *n/3* (64). In an asymmetric tree, tips tend to coalesce with branches deeper in the tree, and there are fewer cherries than expected. Using simulations of differing risk of transmission, Frost and Volz *et al*. (17) demonstrated that higher infectiousness resulted in more asymmetric trees (not observed here), though dependent to an extent on the sampling fraction (also not observed here). As dynamic risk groups were not investigated, we sought to explore whether the proportion of cherries (relative to the number of tips) for a given cluster could aid in the resolution of epidemic dynamics for a given cluster.

### B. Feature description & pre-processing

Based on different raw data formats, we performed corresponding data preprocessing steps. Specifically, for the categorical feature *LTTShape* with four categories in its raw format, i.e., concave, concave_convex, convex, and convex_concave, we utilized a one-hot vector for encoding it. For the continuous feature *gamma* but with infinity value (denoted by *inf*), we selected to use quantile discretization to split it into four groups, i.e., (−4391.276,−16.903], (−16.903,−15.72], (−15.72,−12.722], and (–*12.722,inf*).Then one-hot encoding with four dimensions was utilized to represent different groups. The other continuous variables with limited data ranges including Oster, PD, AbsGrowthRate, FractionTimeGrowth, *R_max_*, Cherries, BLD and *R*_0_ were simply processed with a z-score normalization where each feature was normalized to the zero mean and unit variance. For the two edge features, we first applied inverse hyperbolic sine (short for ArcSinh) transformation, which can approximate the natural logarithm of the raw values and retain zero-valued observations, then standardize them with z-score normalization. Table 1 and 2 respectively provide the data statistics for raw node and edge features.

**Table 1.** Summary statistics for generalized tree statistic features.

| Features | Descriptions | Min | Max | Mean | Std |
| --- | --- | --- | --- | --- | --- |
| *Sampling Fraction* | Fraction of total infected individuals sampled from a cluster | $4.072 \times 10^{-2}$ | 1.000 | $1.455 \times 10^{-1}$ | $2.288 \times 10^{-1}$ |
| *Cluster Size* | Total number of infected individuals in a cluster | 4.000 | $1.103 \times 10^3$ | $5.570 \times 10^2$ | $2.113 \times 10^2$ |
| *Time Span* | Time range from earliest sampled individual to most recently sampled individual within a cluster | 2.000 | $1.540 \times 10^2$ | $8.353 \times 10^1$ | $2.631 \times 10^1$ |
| $\gamma$ | Pybus's $\gamma$ (60) | $-4.391 \times 10^3$ | *inf* | *inf* | *NA* |
| *Oster* | Estimated transmission rate (per 100 infected individuals per time unit) (16)<sup>1</sup> | 2.178 | $1.860 \times 10^2$ | $9.456 \times 10^1$ | $3.244 \times 10^1$ |
| *PD* | Phylogenetic diversity (sum of branch lengths scaled in genetic distance) | 5.000 | $2.276 \times 10^4$ | $1.361 \times 10^4$ | $5.560 \times 10^3$ |
| $R_0$ | Basic reproductive number | $2.429 \times 10^{-4}$ | 8.805 | 1.884 | $6.433 \times 10^{-1}$ |
| *Abs Growth Rate* | Average growth rate for estimated $N_e$<sup>2</sup> | $8.178 \times 10^{-9}$ | $3.186 \times 10^{15}$ | $4.268 \times 10^9$ | $2.721 \times 10^{12}$ |
| *Fraction Time Growth* | Fraction of cluster time span (includes TMRCA<sup>3</sup>) spent in growth phase | 0.000 | 1.000 | $3.035 \times 10^{-1}$ | $2.812 \times 10^{-1}$ |
| $R_{max}$ | Maximum rate of growth in estimated $N_e$<sup>2</sup> | $-5.141 \times 10^7$ | $6.486 \times 10^6$ | $5.503 \times 10^3$ | $7.642 \times 10^4$ |
| *LTT Shape* | Shape of LTT<sup>4</sup> | *NA* | *NA* | *NA* | *NA* |
| *Cherries* | Number of cherries per total number of taxa in cluster | $8.333 \times 10^{-2}$ | $5.000 \times 10^{-1}$ | $3.265 \times 10^{-1}$ | $2.411 \times 10^{-2}$ |
| *BLD* | Branch length difference | $-8.450 \times 10^1$ | $1.300 \times 10^1$ | 4.023 | 3.826 |

<sup>1</sup> Originally proposed for HIV-infected individuals
<sup>2</sup> Effective population size
<sup>3</sup> Time of the most recent common ancestor
<sup>4</sup> Lineages through time (LTT) shape is classified as concave, convex, or a mixture of convex and concave, representing a non-numeric feature with no descriptive statistic information.

**Table 2.** Summary statistics for edge features.

| Features | Descriptions | Min | Max | Mean | Std |
| --- | --- | --- | --- | --- | --- |
| *Weight1* | Time (days) | 0.000 | $1.060 \times 10^2$ | $1.159 \times 10^1$ | 9.450 |
| *Weight2* | Genetic distance (substitutions/site) | 0.000 | $8.971 \times 10^{-2}$ | $1.043 \times 10^{-2}$ | $8.549 \times 10^{-3}$ |

### C. Feature distribution

Fig. 1 and 2 illustrate the distributions of the raw and normalized tree statistic features on both training (including validation) and testing datasets. The three ground truth cluster characteristics, i.e. Sampling Fraction, Cluster Size and Time Span, are also included. In order to provide a better visualization, samples with *Abs Growth Rate* > 100 are not shown in Fig. 1 and 2. Fig. 3 and 4 respectively exhibit the Spearman’s rank and Pearson correlation between tree shape statistic features (including the three ground truth cluster characteristics). Fig. 5 illustrates the distribution of the two edge features. Fig. 6 exhibits the statistic information of the three dynamic transmission cluster types in training and test datasets.

**Fig. 1.**
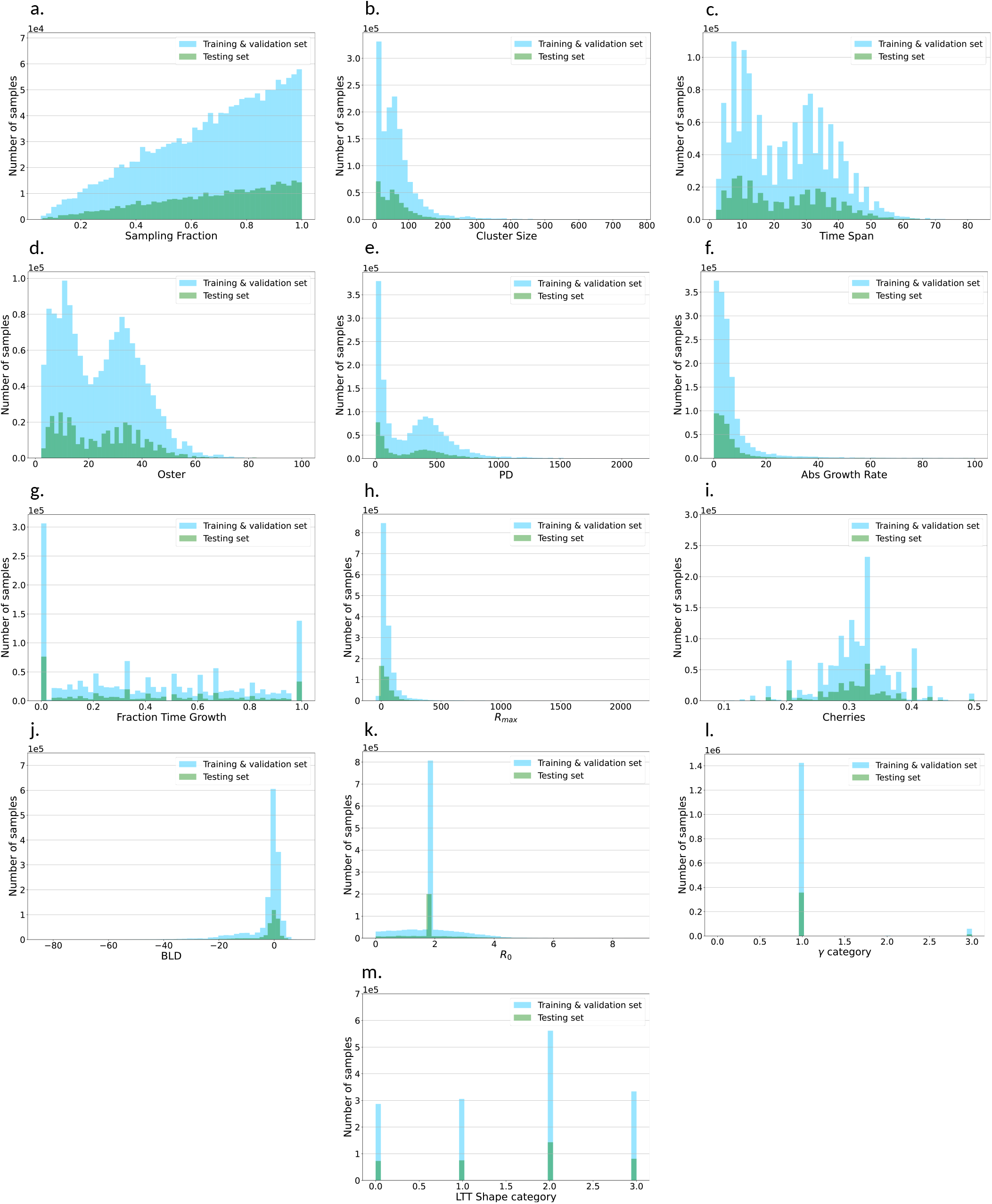
Distribution of the raw tree shape statistics with numerical values.

**Fig. 2.**
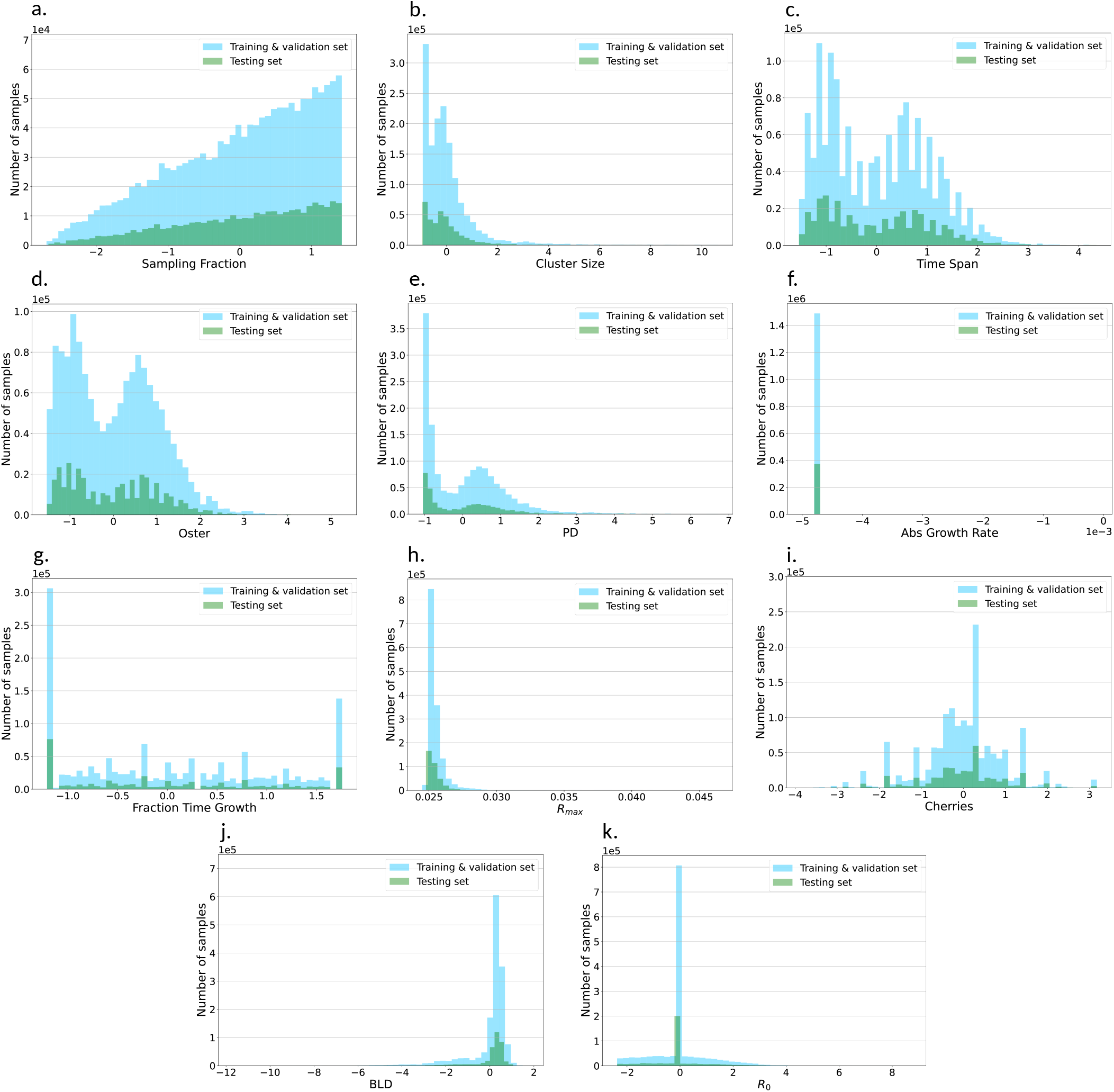
Distribution of the normalized tree shape statistics with numerical values.

**Fig. 3.**
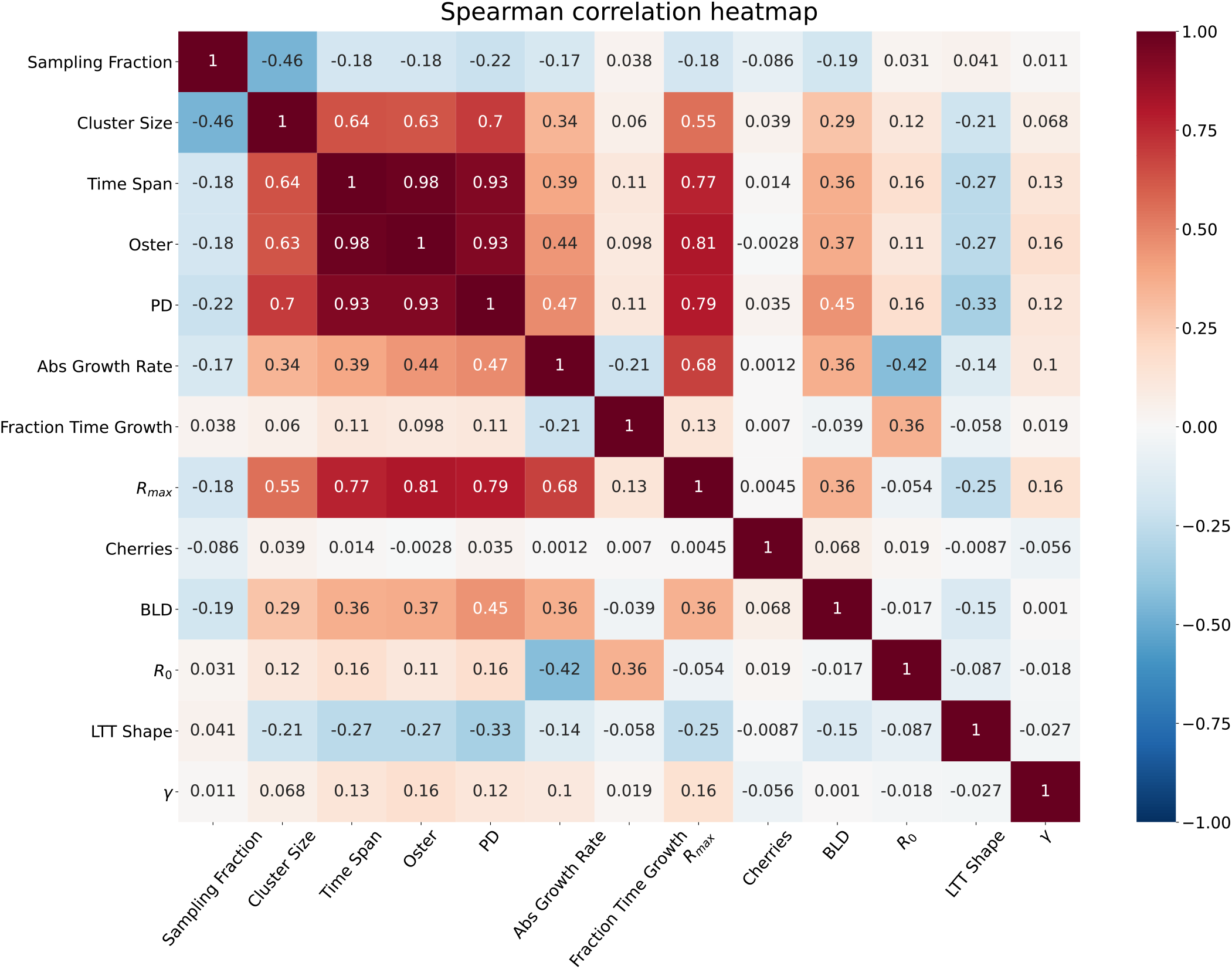
Spearman’s rank correlation between the ten generalized tree shape statistics and three ground truth cluster characteristics, including sampling fraction, size and time span of the cluster.

**Fig. 4.**
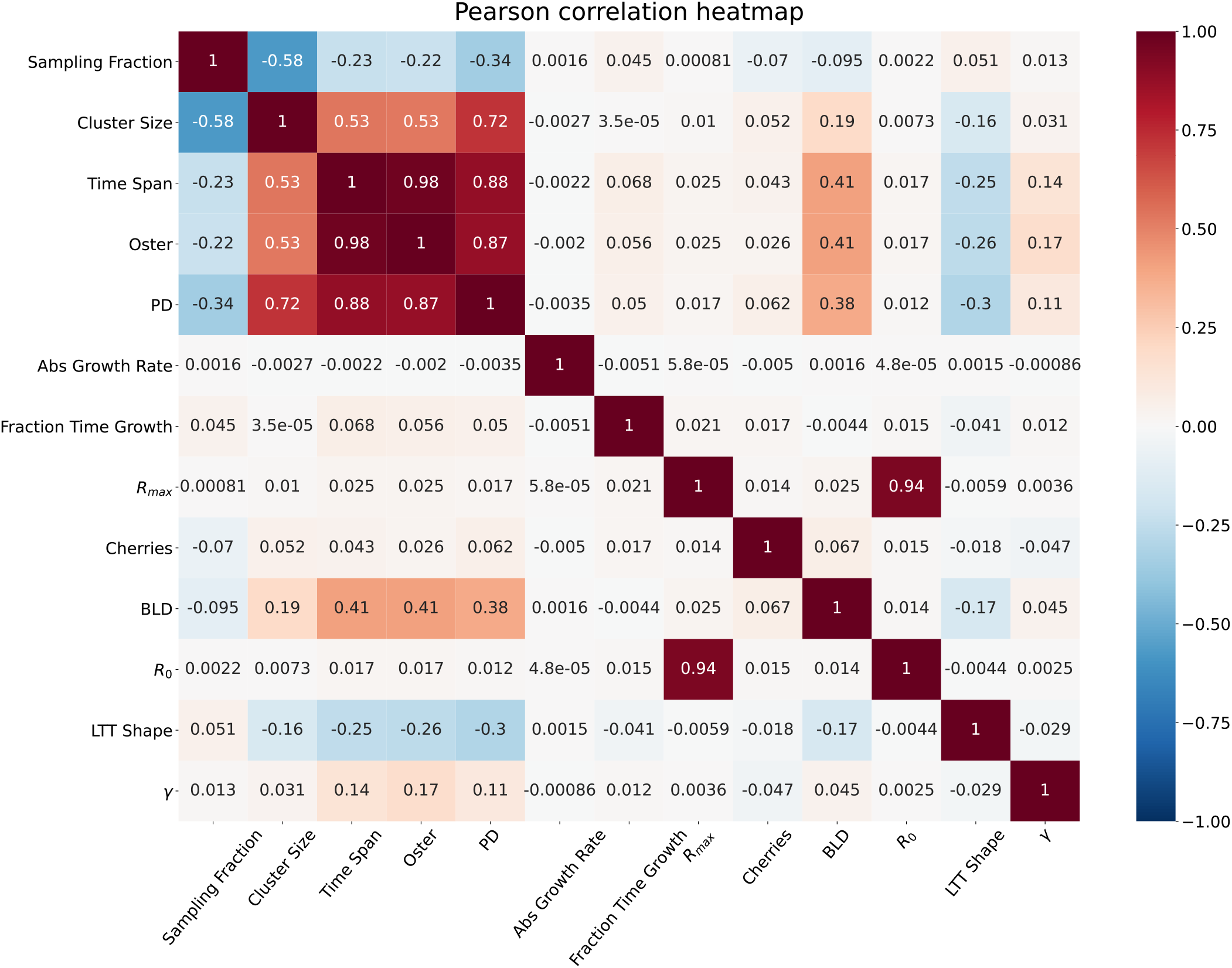
Pearson correlation between the ten generalized tree shape statistics and three ground truth cluster characteristics, including sampling fraction, size and time span of the cluster.

**Fig. 5.**
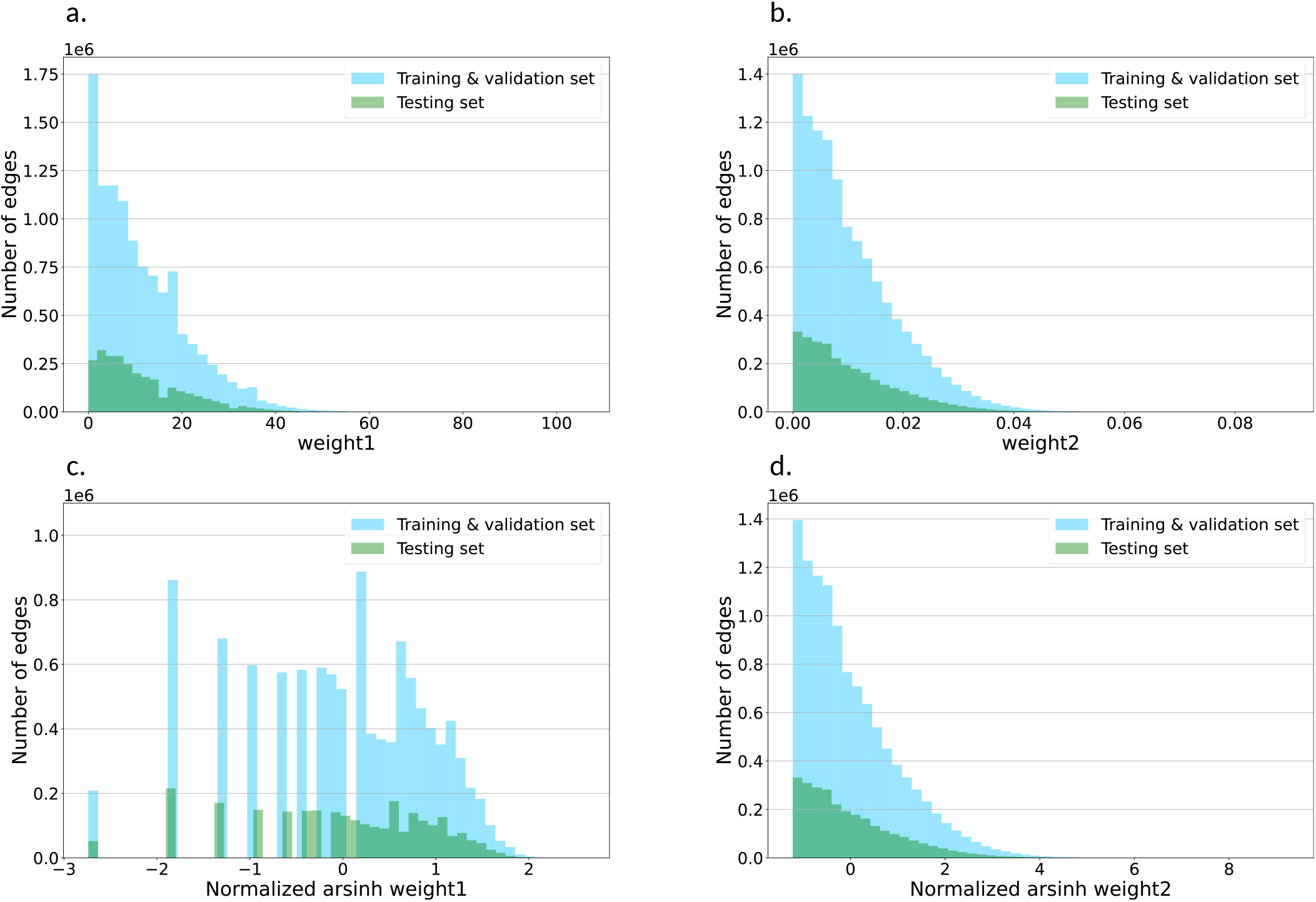
Distribution of the edge features. **a** and **b** are distributions of the raw edge features; weight1 and weight2. **c** and **d** are distributions of the edge features processed by an ArcSinh transformation and a z-score normalization.

**Fig. 6.**
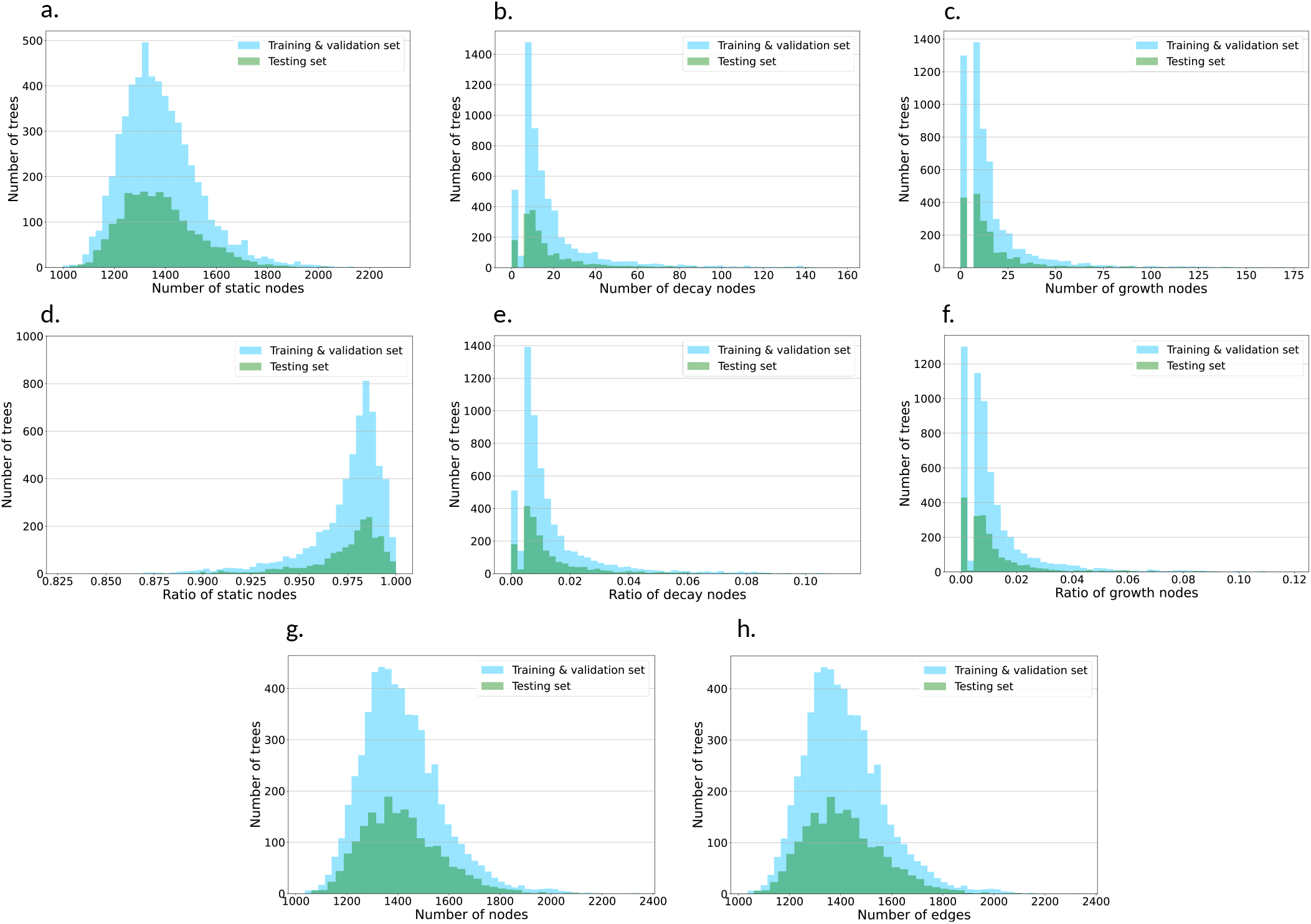
Statistic information for three different dynamic transmission clusters types, i.e. static, decay and growth. **a, b and c** are the histogram of number of static, decay and growth nodes among the trees, and plot d.-f. show the distribution of classes’ ratio. Static nodes show the dominate portion of our dataset. Plot g. and f. show histogram of the number of nodes and edges among trees.

### D. Additional experimental results

Fig. 7 illustrates the performance comparison of all the benchmarking algorithms using the averaged receiver operator characteristic curves, confusion matrices, and permutation feature importance results. Fig. 8 shows the permutation feature importance results of our *DeepDynaTree-PDGLSTM*, and the absolute reduction of balanced accuracy on the testing dataset was used as the measurement. Fig. 9 provides an illustration on how to construct a dual graph from a phlogenetic tree.

**Fig. 7.**
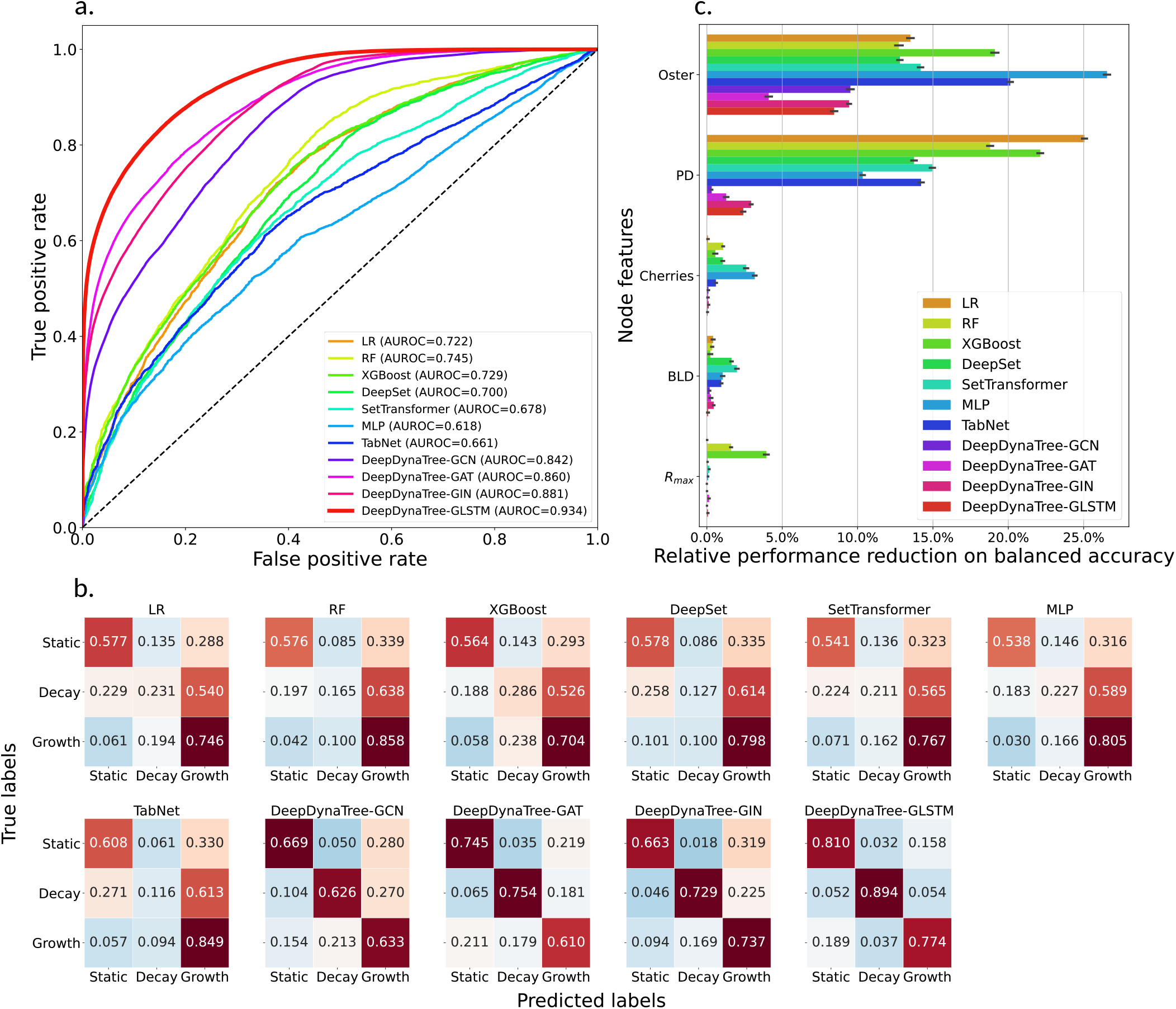
Figurative performance comparison of all classifiers and permutation feature importance results. **a,** Comparison of various classifiers, shown on a equally averaged receiver operator characteristic curve, with AUC indicated for each classifier. **b,** Comparison of various classifiers, shown on confusion matrices, and elements are row-wise normalized by class support size. **c,** Relative permutation feature importance results measured by the balanced accuracy. x-axis represents the relative model performance reduction in percentage compared to the non-permutated model and y-axis represents the top-5 most important generalized tree shape statistics features. The features are ranked based on the average relative reduction over all the models, and their relative reductions’ mean and standard deviation were measured over 50 runs.

**Fig. 8.**
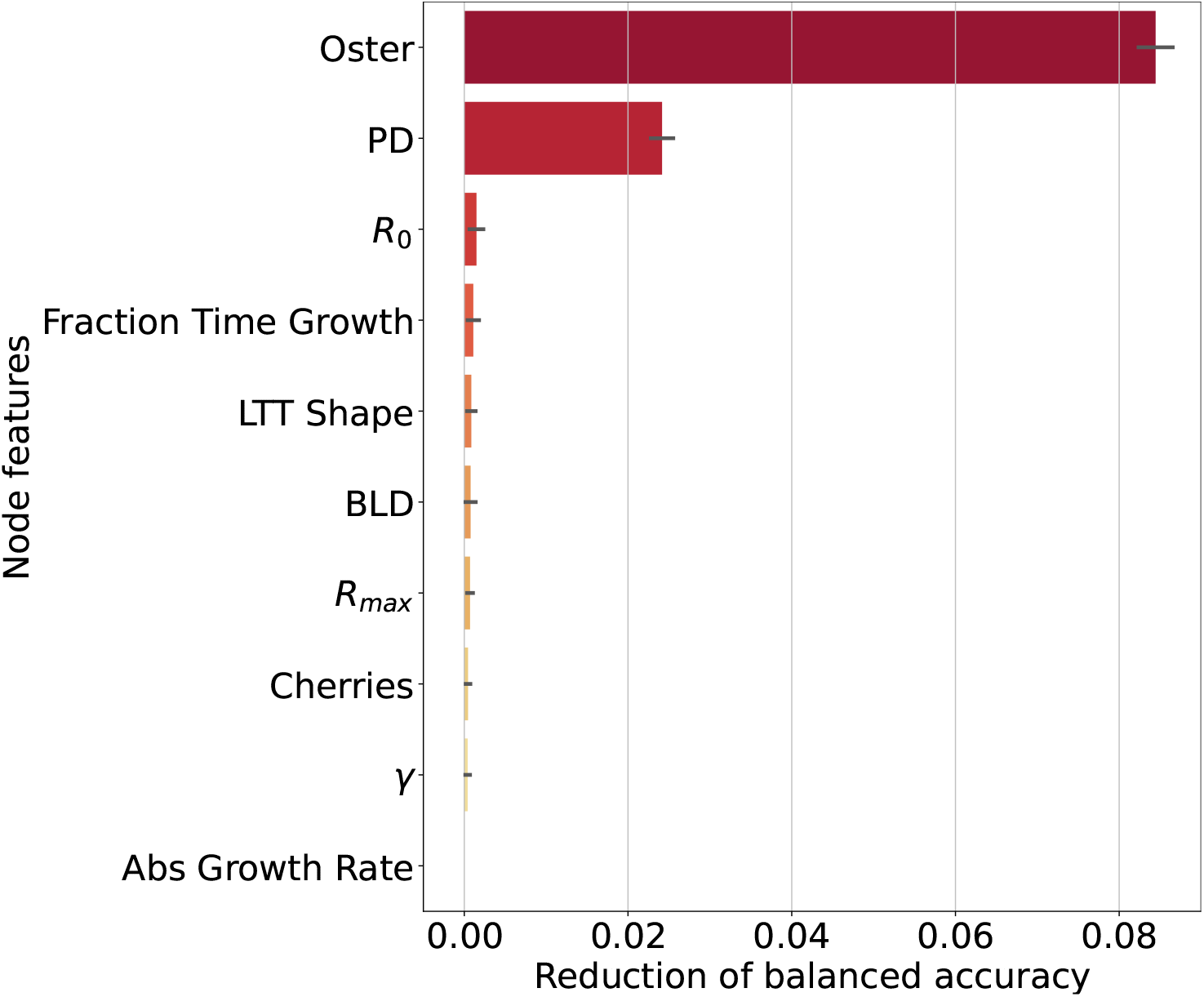
Permutation feature importance results for *DeepDynaTree-PDGLSTM*. Features are ranked by the permutation importance scores. The mean and variance of each feature’s permutation importance were measured over 50 runs.

**Fig. 9.**
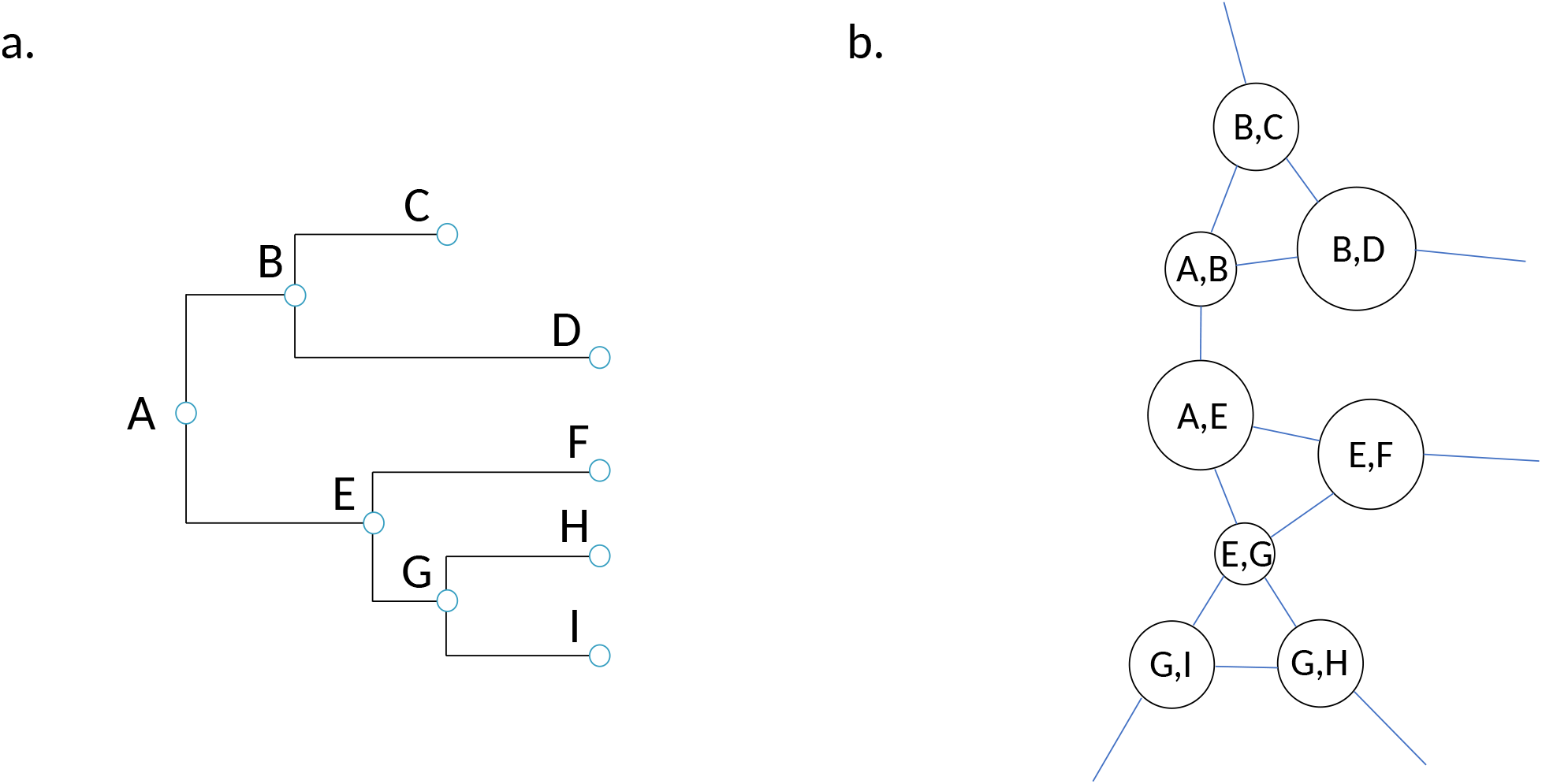
An illustration of a phylogenetic tree and its corresponding dual graph. **a,** Each branch point in the phylogenetic tree represents a transmission cluster, and the various branching lengths represent different genetic distances or evolutionary times separating individual nodes. **b,** The vertices and edges in the dual graph correspond to the branches and nodes of the original phylogenetic tree respectively. The size of the node cycle in the dual graph is proportional to the corresponding branch length in the phylogenetic tree.

